# Dynamic Resting-State Network Markers of Disruptive Behavior Problems in Youth

**DOI:** 10.1101/2025.05.15.654366

**Authors:** Heather M. Shappell, Zhiyuan Liu, Mohammadreza Khodaei, George He, Dylan G. Gee, Martin A. Lindquist, Denis G. Sukhodolsky, Gregory McCarthy, Karim Ibrahim

## Abstract

**Background:** Childhood disruptive behavior problems are linked to aberrant integrity within large-scale cognitive control networks. However, it is unclear if transitory or dynamic variation in the functional brain architecture is a marker of disruptive behavior problems. The current study tested whether functional connectivity across dynamic networks is distinctly associated with the transdiagnostic symptom domain of disruptive behavior problems in children.

**Methods:** Participants were aged 9-10 years from the Adolescent Brain Cognitive Development (ABCD) Study, who completed resting-state fMRI (N=877). We employed a dynamic connectivity approach leveraging a hidden semi-Markov model (HSMM) to identify transient properties of brain networks and states. Models estimated the time spent in each state (occupancy time) and the number of consecutive timepoints in a state (dwell time) for each participant. Linear regression models were utilized to identify distinct associations between dynamic properties (occupancy and sojourn times) and severity of disruptive behavior problems, accounting for other commonly co-occurring symptoms.

**Results:** Dynamic network markers of disruptive behavior problems included increased time in network states characterized by globally aberrant connectivity patterns in circuitry involved in cognitive control including frontoparietal and dorsal attention networks. Replication of findings was found in a held-out sample of resting-state fMRI runs in which greater severity of disruptive behavior problems was uniquely linked to greater occupancy time in similarly characterized brain states.

**Conclusion:** Transdiagnostic, dynamic resting-state markers of disruptive behavior problems in youth may assist in the development of brain-based biomarkers for monitoring treatment outcomes, assessing circuit target engagement and informing clinical decisions.

## Introduction

Childhood disruptive behavior disorders (DBD), including Oppositional Defiant Disorder and Conduct Disorder, form one of the most prevalent classes of childhood-onset psychiatric disorders worldwide (1, 2). Childhood DBD are diagnosed based on the presence of symptoms including irritability/anger, aggression, noncompliance, and/or antisocial behavior (2). Additionally, childhood disruptive behavior is also recognized as a transdiagnostic dimension (3) present across various psychiatric disorders (4, 5). DBD are marked by aberrant connectivity across networks implicated in the top-down control of emotion, or emotion regulation, and cognitive control more broadly (6–15). For instance, recent work indicates altered connectivity across functional networks spanning cognitive control, social perception, and emotion processing circuitry linked to disruptive behavior severity in a transdiagnostic sample of youth (10). Thus, aberrant connectivity across cognitive control networks supporting executive functioning, inhibitory control, cognitive flexibility, and emotion regulation may be a shared mechanism underlying childhood disruptive behavior (8, 9, 16–18). Previous studies of DBD have all used “static” connectivity approaches, which derive an average network connectivity metric for the length of time an individual is in a scanner. To our knowledge, no study to date has examined whether alterations in the moment-to-moment fluctuations in functional connectivity (i.e., dynamic functional connectivity) are linked to disruptive behavior severity in youth. As such, it remains largely unknown whether dysfunction in the time-varying properties of cognitive control networks is implicated in disruptive behavior severity and if such effects reflect trait (static) versus state (dynamic) variation in the functional brain architecture.

Recently developed methods assess changes in dynamic functional connectivity and implement data-driven approaches to identify distinct patterns or “states”, which can advance understanding of network organization associated with symptom dimensions in youth (19–22). Disruptions in the dynamic properties of functional networks, including cognitive control circuits, have been linked with symptom severity, primarily in adults (23–31). For instance, evidence links dysfunction in the amount of time spent in network configurations (dwell time or occupancy time) with schizophrenia, depressive disorders, and autism spectrum disorder (ASD) in adults (23, 25, 26, 29, 30, 32): that is, patient groups may spend greater time in brain states characterized by reduced connectivity compared to unaffected controls. For instance, Reinen et al. (32) reported that longer dwell time in hypo-connected, cognitive control networks coupled with fewer transitions across states was evident among adults with schizophrenia. Recent work also shows that disruptions in dwell time and state transitions in large-scale cortical networks predict ADHD in children relative to controls (31). Specifically, children with ADHD may spend less time in and switch more quickly out of hyperconnected states involving cognitive control circuitry, including the default mode and task-relevant networks, compared to unaffected controls (31). Thus, data-driven, dynamic connectivity approaches complement information gained from static connectivity approaches and may hold promise in identifying altered functional interactions among networks in the human connectome that are associated with childhood disruptive behavior, facilitating the identification of robust brain-based biomarkers (33). For instance, brain biomarkers derived from static and dynamic connectivity can facilitate precision treatments by serving as ‘neural fingerprints’ to identify patients likely to benefit from a particular intervention and inform circuit location for neuromodulation. While previous research has implicated a broader network dysfunction in childhood DBD (10), dynamic connectivity approaches offer significant advancements by providing a more detailed understanding. These approaches allow for the examination of moment-to-moment reconfigurations of cognitive control networks, independent of the implicit assumptions associated with static connectivity analyses. Importantly, this nuanced evaluation could yield precise insights into the underlying dysconnectivity of disruptive behavior problems in youth.

Here, we employed a dynamic connectivity approach that leverages a hidden semi-Markov model (HSMM) to identify transient properties of brain networks associated with disruptive behavior severity (31, 34). HSMMs are a novel approach to dynamic connectivity analysis that have been shown to complement the more traditional sliding window analysis approach (34, 35). The HSMM approach has several advantages including handling large amounts of complex data, and unlike the sliding window analysis, identifying rapid changes in dynamic connectivity. HSMM approaches also have the major advantage, in comparison with standard hidden Markov models, of explicitly modeling and estimating dwell time distributions. We utilized HSMMs to assess transitions, dwell times, and total occupancy times across distinct functional connectivity states in a transdiagnostic sample of children with varying levels of disruptive behavior. In the current study, we applied HSMMs to resting-state fMRI data from the Adolescent Brain Cognitive Development (ABCD) Study, (36, 37). HSMMs were used to identify distinct brain states and brain-behavior associations between a continuous measure of disruptive behavior—measured using the Child Behavior Checklist (CBCL) Externalizing Behavior Problems subscale (38)—and dynamic functional connectivity across large-scale resting-state networks in a transdiagnostic sample of children. Based on previous static connectivity work implicating aberrant cognitive control connectivity in childhood disruptive behavior (6, 8, 11, 39), we hypothesized that altered dynamic connectivity among cognitive control networks would be associated with the severity of disruptive behavior problems.

The present study extends prior work in at least three ways. First, prior studies have focused on static functional connectivity correlates of disruptive behavior severity. Recent work by Shappell et al. (31) provided critical insight into the functional relevance of dynamic network states to ADHD in children using HSMM. The present study complements this work by evaluating, for the first time, the relevance of dynamic connectivity states to disruptive behavior severity. Second, we extend prior work by testing a transdiagnostic and diverse sample of children with varied demographic backgrounds from the ABCD Study (40). Third, we apply an HSMM approach, identical to Shappell (31, 34), to elucidate brain states unique to disruptive behavior problems and facilitating comparison between studies and age groups.

## Materials and Methods

### Participants

We analyzed a subset of participants from the ABCD Study (36) (https://abcdstudy.org/; baseline assessment [release 2.0.1]). The ABCD Study is a large longitudinal study with over 11,500 children that aims to comprehensively characterize psychological and neurobiological development from early adolescence to early adulthood (36). Details of the ABCD Study are described in prior work (36, 37) and elsewhere for recruitment (41), neurocognitive batteries (42), and imaging protocols (37). As the combination of HSMM brain state estimation analyses with independent component analysis are computationally intensive, fitting all models to the full sample during model development and testing would have been incredibly time intensive. Therefore, we randomly selected a subset of 877 (420 females) individuals ages 9 to 10 years with varying levels or severities of problem behaviors or symptoms with 4 runs of baseline resting-state fMRI that met the ABCD Study recommended inclusion criterion for structural MRI and resting-state fMRI quality control. In order to test the replication of associations between brain network dynamics and disruptive behavior severity, primary analyses were conducted on 2 runs of resting-state fMRI data (discovery dataset). Replication analyses were performed on the remaining held-out 2 runs of resting-state fMRI data. The ABCD’s Data Analysis, Informatics and Resource Center (DAIRC) (37) provides a dichotomously coded variable of the recommended sample for each imaging modality. Additional detail is provided in the **Supplemental Methods**. Sample characteristics are shown in **Table 1**.

**Table 1.**
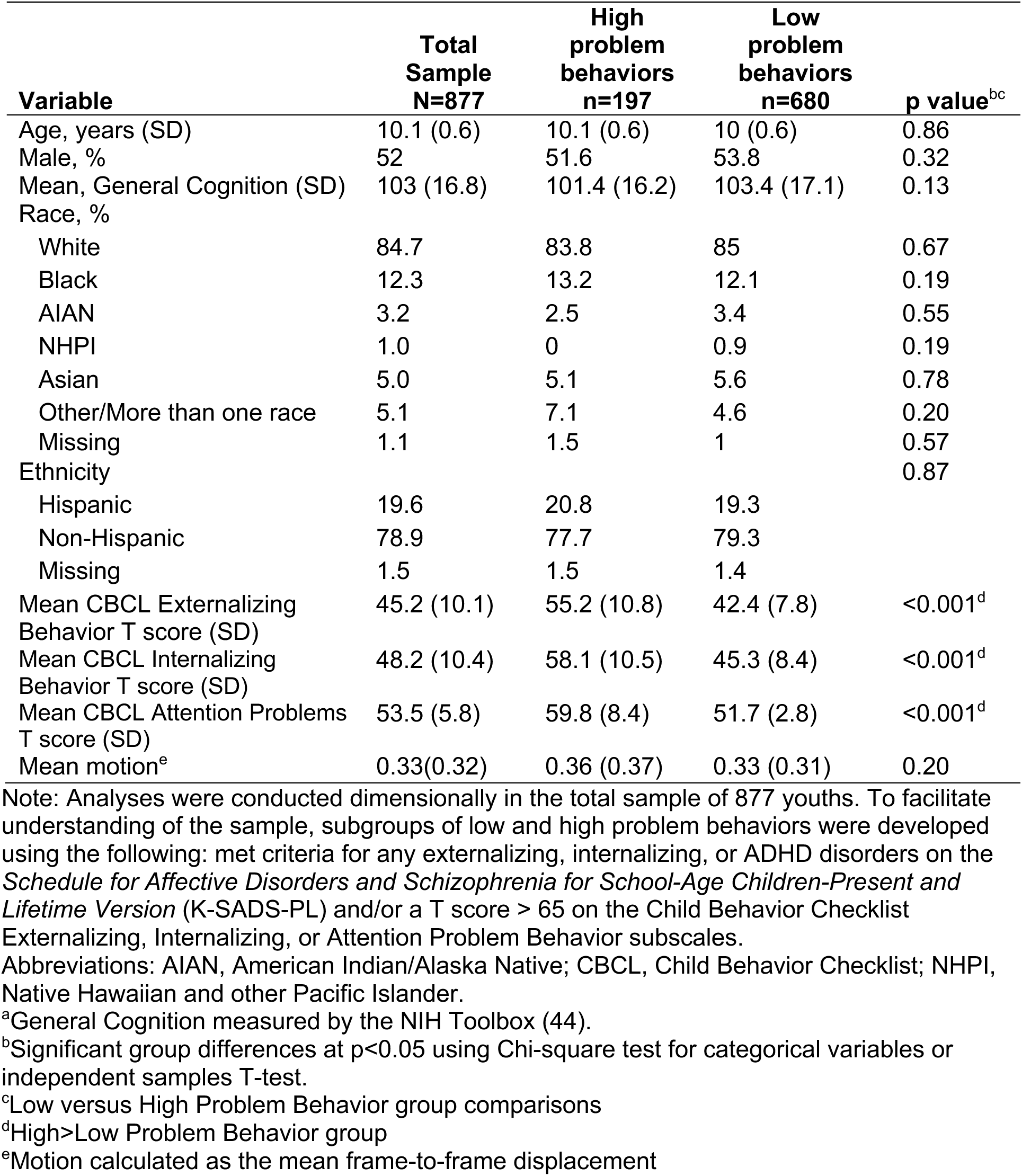
Participant demographics and clinical characteristics.

### Behavioral Measures

We used a continuous measure of symptom domains, which was selected to allow comparison with prior imaging studies using the ABCD dataset (10, 13, 43). The parent-rated Child Behavior Checklist (CBCL) (38), which is a well-established measure of child psychopathology, provides broadband measures of transdiagnostic symptom domains. The *CBCL Externalizing Behavior Problems* scale assesses disruptive behavior problems related to verbal and physical aggression, conduct problems (e.g., attacks, setting fires, running away, rule-breaking, truancy), anger/irritability, and noncompliance. The *CBCL Internalizing Problems* scale includes items assessing somatic complaints, social withdrawal, and anxiety/depressed symptoms. The *CBCL Attention Problems* scale assesses inattention, hyperactivity, and impulsivity symptoms. Participants also completed cognitive assessments including the NIH Toolbox Cognition battery (44). The NIH Toolbox Cognition total composite score was used in analyses to account for cognitive performance.

### Imaging Acquisition and Processing

FreeSurfer v6.0 (http://surfer.nmr.mgh.harvard.edu) processing stream was applied to each participant’s high-resolution T1-weighted scan, which has been previously described (45, 46). Minimally preprocessed resting-state fMRI data were processed using Functional MRI of the Brain (FMRIB) Software Library (FSL Version 4.1.6; FMRIB, Oxford, United Kingdom) (47, 48) and denoised using the AROMA package (49) as described in our previous work (9, 10). Participants completed at least 8 minutes of eyes-open resting-state. For details on the ABCD imaging protocol, see Hagler et al. (37). More detail regarding preprocessing and denoising is provided in the **Supplemental Methods**.

### Independent Component Analysis

Large-scale intrinsic networks were identified at the group level based on a data-driven approach using FSL’s Multivariate Exploratory Linear Optimized Decomposition into Independent Components (50) (MELODIC) tool (v3.15). A multisession temporal concatenation approach was used to generate a parcellation of 50 spatiotemporal components covering both the cortex and subcortex. ICA components were mapped to Yeo intrinsic networks as in previous work (51). Following removal of components determined to be motion or artifact, 33 ICA components were included in the final analysis. Full details are provided in the **Supplemental Methods**.

### Hidden semi-Markov modeling (HSMM)

We fit an HSMM on the ICA time series data of our final sample of study participants. A detailed description of the HSMM can be found in our prior work (31, 34), but we provide a summary here. We denote the ICA time-series data for each participant by *Y_i_*_1_*, …, Y_iT_*, where each *p*−dimensional vector *Y_it_* contains the BOLD measurements of the *p ICA* components at the *t^th^* timepoint for the *i^th^* participant. Now suppose a unique unobservable (or hidden) brain network state gives rise to each *Y_it_*. We represent this hidden network state at a particular observation time by *S_it_*, where *S_it_* takes discrete values (i.e., *S_it_* ∈ 1*…K*). We assume that each *Y_it_* follows a multivariate Gaussian distribution *Y_it_* ∼ N (*µ_s_,* Σ*_s_*), where the mean and covariance are dependent on the current (unknown) network state, for any *S* ∈ 1*, …, K*. Thus, each network state has its own set of mean activations across our ICA components, as well as its own covariance structure between components. We convert the covariance matrix for each network state to a correlation matrix, which represents the weighted brain network for each state. The HSMM also contains parameters that estimate a) transition probabilities between network states and b) dwell time distributions for each state. See **Supplementary Methods** for the complete data log-likelihood of the HSMM.

To ensure that brain states represent subgroups of youths with low and high levels of symptoms across psychopathology domains, we fit states separately on each subgroup. The number of states one can fit must be specified a-priori and is heavily dependent on the number of participants, number of timepoints, and number of ICA components included in the analysis. Given our sample size, we found that six states produced stable parameter estimates over multiple fittings of the model. Therefore, 6 states were fitted separately for the subgroup with high symptomatology (based on a cutoff of T score > 65 on the CBCL Internalizing, Externalizing, or Attention scales and/or meets DSM-5 criteria for internalizing disorders including anxiety and depressive disorders, DBD, or ADHD) and for the subgroup with low symptomatology (T score < 55 on the same CBCL scales and/or no current DSM-5 criteria met for any diagnosis). Next, the model was fit using the combined 12 states (6 states from low and 6 states from high levels of problem behaviors subgroups) on our entire sample of 877 participants. Fixing the 12 states estimated from each subgroup separately ensures representation of brain states along a continuum of psychopathology domains. All HSMM parameters were estimated utilizing the *mhsmm* R package (52).

Each participant’s most probable sequence of true network states was estimated using the Viterbi Algorithm (54). The Viterbi algorithm takes an individual’s time-series data and finds which of the 12 states estimated from the entire data set that the data is suggesting they are in at each timepoint. Metrics were extracted from these state sequences (such as total time spent in each state), for each subject, which were then implemented in linear regression models to test for associations with externalizing behavior severity (see next section).

### Statistical Inference

To examine the relationship between externalizing behavior severity and state occupancy time (dependent variable), we conducted linear regression models in R. The occupancy time for a particular state represents the total number of timepoints during the fMRI scan that the individual spent in the state. Regression models accounted for potential confounds and included age, sex as a dichotomously coded variable, and cognitive performance. To ensure specificity of findings for disruptive behavior problems, models also accounted for severity of attention and internalizing problems (using the CBCL Attention and Internalizing Problems subscale scores, respectively). Regression models were fit to each of the 12 states. A false discovery-rate (FDR) correction using the Benjamini-Hochberg algorithm in R (p.adjust function) was applied across all models. Follow-up analyses were conducted to account for motion. Topology of states and intrinsic connectivity were visualized using the R package ‘igraph’ and the Fruchterman-Reingold algorithm layout (55), in which highly connected nodes are placed near each other.

### Topological Characteristics of Brain States

To further understand the topological characteristics of brain states so that we may better characterize each brain network state, we implemented graph theory to derive metrics of degree, betweenness centrality, local and global efficiency, and clustering coefficients. Undirected, weighted graphs were derived from state covariance matrices estimated from the HSMM. Each covariance matrix was converted to a correlation matrix and all negative correlations were set to 0. We implemented the Brain Connectivity Toolbox (56) (www.brain-connectivity-toolbox.net) to estimate measures of global properties (local and global efficiency) and nodal properties (degree, betweenness centrality, clustering coefficient) for each state.

### Supplemental Follow-Up Analysis of Modularity for Brain States

For any brain states emerging from primary analyses that showed significant and distinct or specific associations with disruptive behavior problems, we conducted follow-up analyses of modularity to further understanding of that state’s characteristics. Specifically, we used a modularity analysis to identify community structures within each state. Modularity analysis estimates communities of nodes that have strong within-group connections while their between-group connections remain weak (57). As a result, information flow is higher within communities compared to between communities. We applied the Newman spectral community detection method to the positive network of each state (58). A gamma value of 1.25 was used to divide the networks into communities. To visualize these modules, we employed circular graphs (59). To prevent overcrowding in the figures, only positive edges, thresholded at 0, that survived a threshold of 0.25 were included.

### Replication Analysis

To assess the robustness of findings across the sample, we tested replication of findings in a held-out sample of two resting-state runs for each subject. That is, the HSMM and linear regression models were repeated using the two held-out runs of resting-state to identify brain states. Large-scale functional networks were derived using the identical approach above in 12 states. Thirty-three independent components were retained and 17 were identified as artifact. Regression models were repeated to test replication of brain-behavior associations. See the **Supplemental Methods** for additional details.

### Data and Code Availability

Data from the ABCD Study are shared on the National Institute of Mental Health Data Archive (NDA) https://nda.nih.gov/. To promote transparency, all code used for analyses is available on GitHub (https://github.com/EmotionNeuroscienceLab).

## Results

### Independent Components Analysis and Network Identification

Large-scale functional networks derived from MELODIC-ICA were mapped onto the resting-state networks defined by Yeo et al.(60) and a subcortical network. In total, 33 independent components were mapped onto 10 large-scale networks (visual, somatomotor, dorsal and ventral attention, limbic, control, temporal parietal, default mode, cerebellar, and subcortical networks) (**Figure 1**).

**Figure 1.**
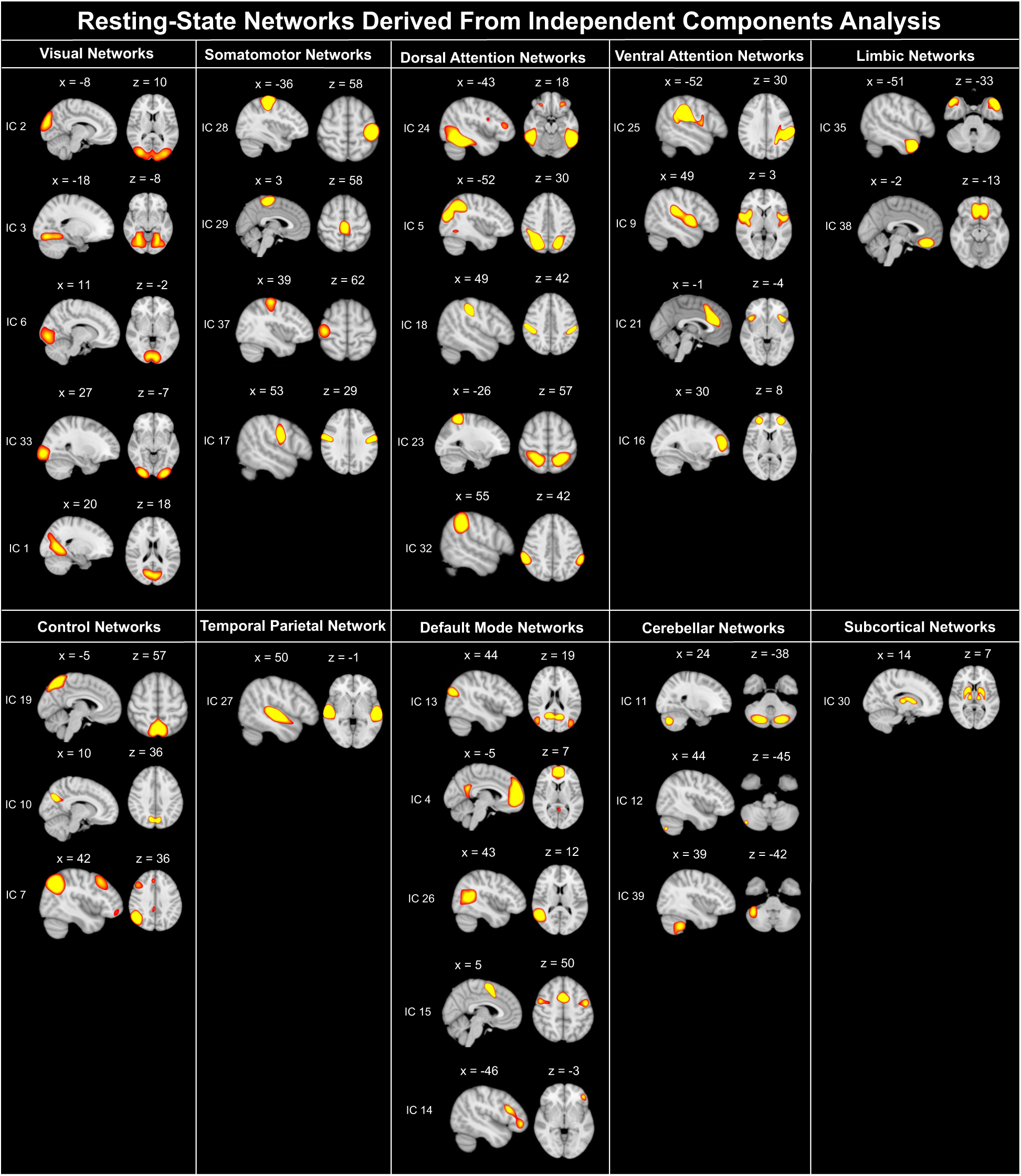
Maps of independent components and grouping into intrinsic connectivity networks. Components were derived using Independent Component Analysis (ICA) implemented using FSL MELODIC. Thirty-three independent components were mapped onto 10 large-scale networks based on the Yeo parcellation.

### Brain State Characterization

Twelve network states were estimated using the HSMM (**Figure 2A**). For visualization purposes, we also provide the thresholded brain state matrices using the top 25% of significant edges (**Figure 2B**). Each state was comprised of multiple large-scale networks, emphasizing the complexity of the functional connectome (61–63). Connectivity was identified within and between networks implicated in cognitive control (frontoparietal, attention), social functioning (default mode, temporal parietal), emotion processing (subcortical, limbic), and sensorimotor functioning (visual, somatomotor, cerebellar).

**Figure 2.**
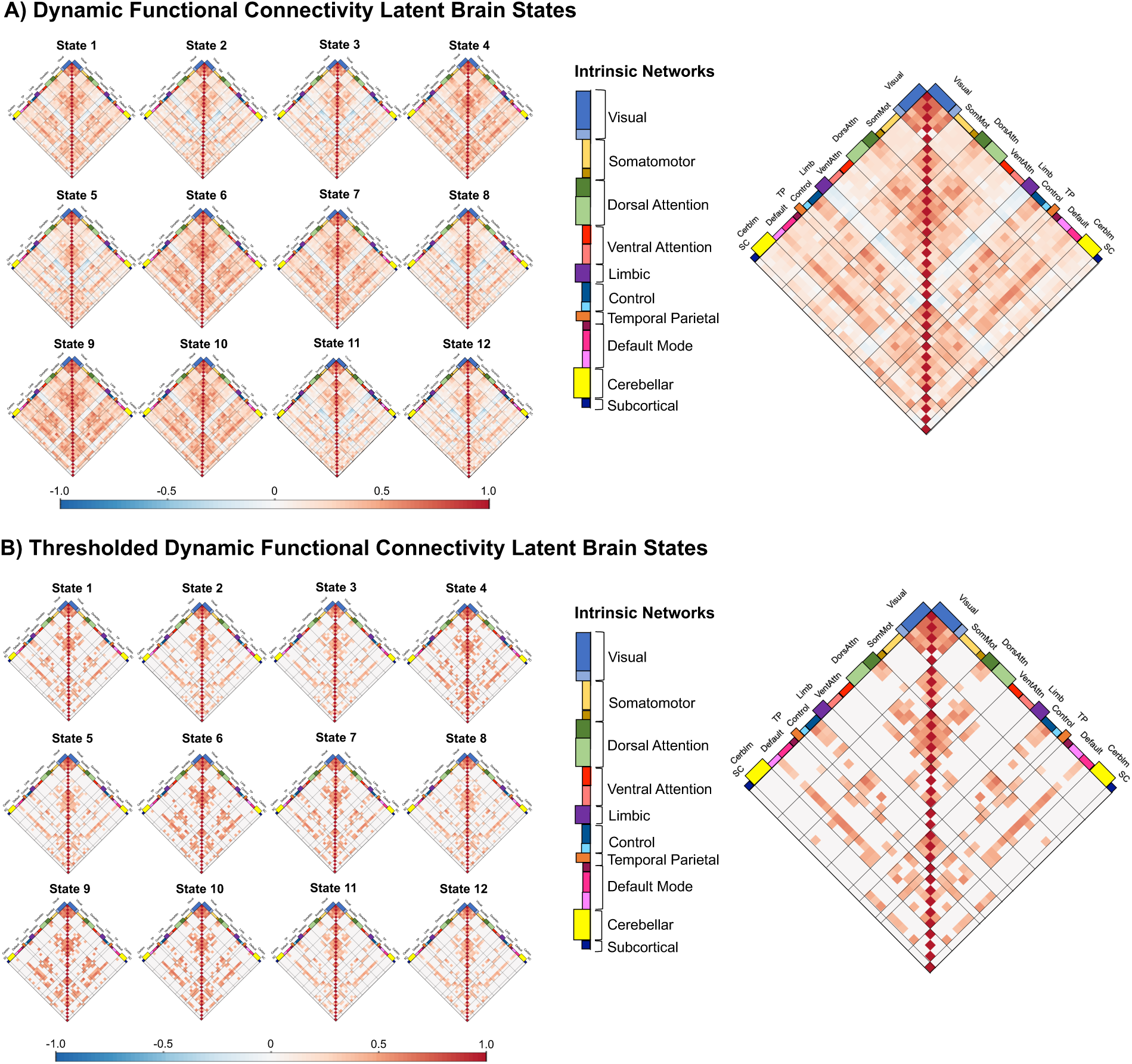
Brain states estimated with HSMM across distributed intrinsic networks. Correlation matrices for states 1 through 12 are shown for each intrinsic network (A) and after thresholding based on the top quartile of significant edges (B). Values reflect Pearson correlations between every region with all other regions for the 12 states. These 12 states are representative of the patterns of functional connectivity in all participants during the resting-state fMRI. Red cells indicate a positive correlation while blue cells indicate a negative correlation (or anticorrelation) between two corresponding ROIs. Abbreviations for intrinsic networks: Cerblm, cerebellar; Control, Frontoparietal control network; Default, default mode network; DorsAttn, Dorsal Attention Network; Limb, limbic; Visual, visual network; SomMat, somatomotor; SC, subcortical; TP, temporo parietal; VentAttn, Ventral Attention Network.

Estimated transition probability matrices were obtained from fitting the models, which showed a pattern of attractor states (32) (i.e., states more likely to be transitioned to) (**Figure 3A-C**). States 10-12, particularly state 10, were more likely to be transitioned to from other states. A more detailed description of the network states associated with disruptive behavior problem severity will be presented in later sections.

**Figure 3.**
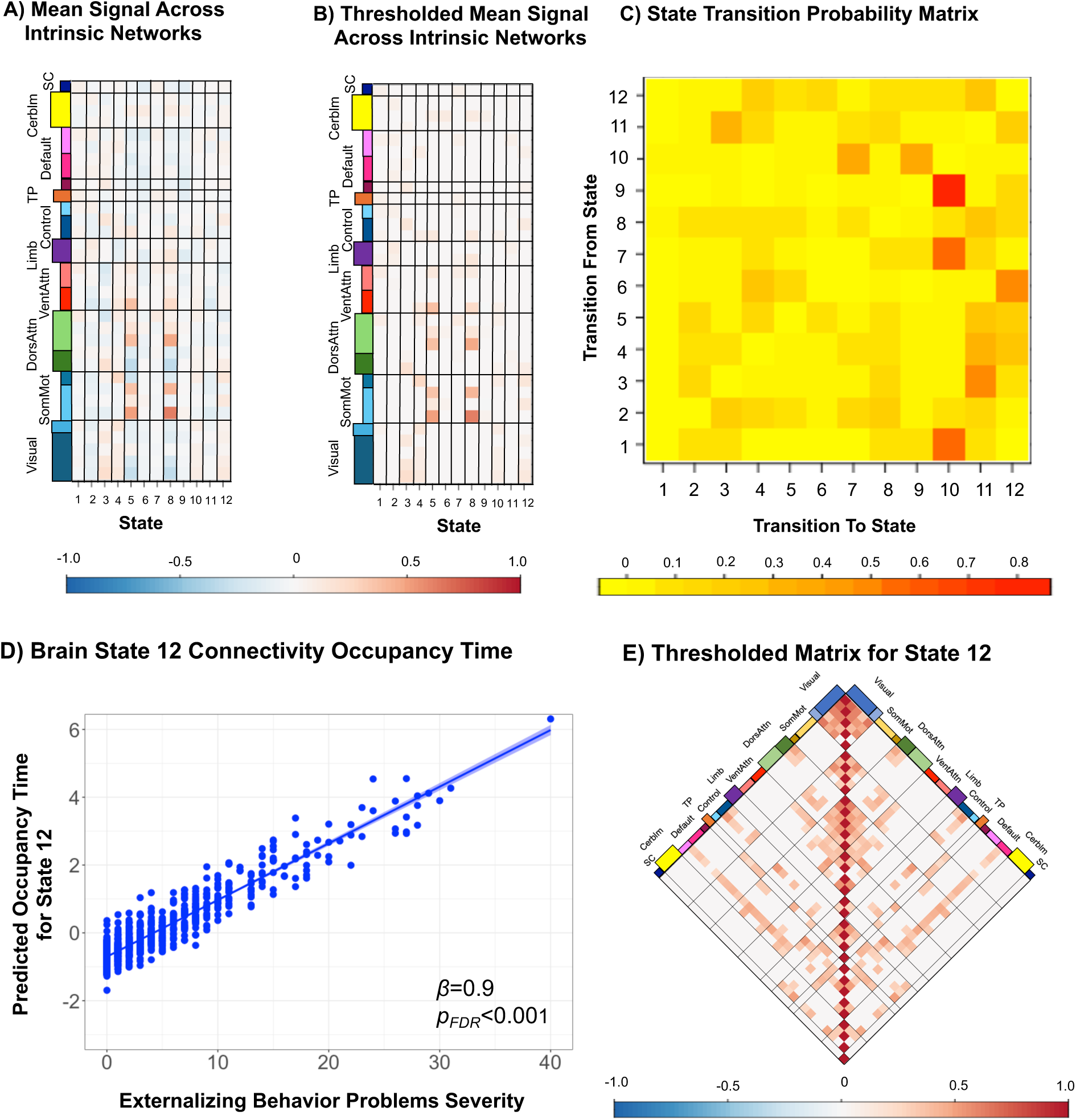
Mean connectivity and transition matrices estimated from HSMM modeling, and occupancy time for state 12 linked to disruptive behavior severity. Mean connectivity is shown for each state (A) and after thresholding based on the top quartile of significant edges (B). Red cells indicate a positive correlation while blue cells indicate a negative correlation (or anticorrelation) between two corresponding ROIs. Transition matrices are also shown for the HSMM model, which indicates probabilities that are conditional on switching to a new state (C). Occupancy time in brain state 12 is distinctly associated with severity of disruptive behavior problems, which is shown in the regression plot. P values are FDR-corrected. See Supplemental Results Table 2 for full regression model parameters.

**Table 2.**
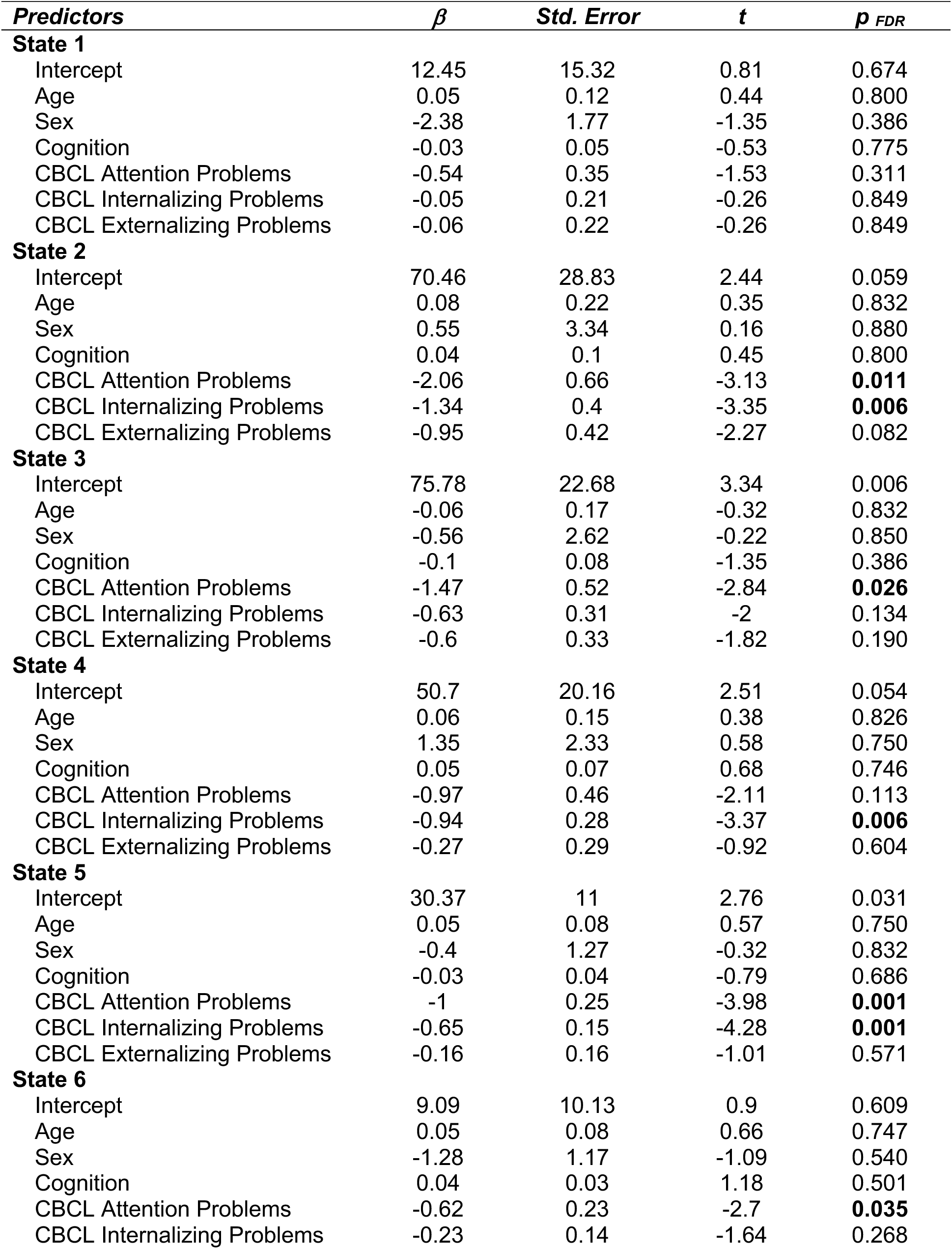
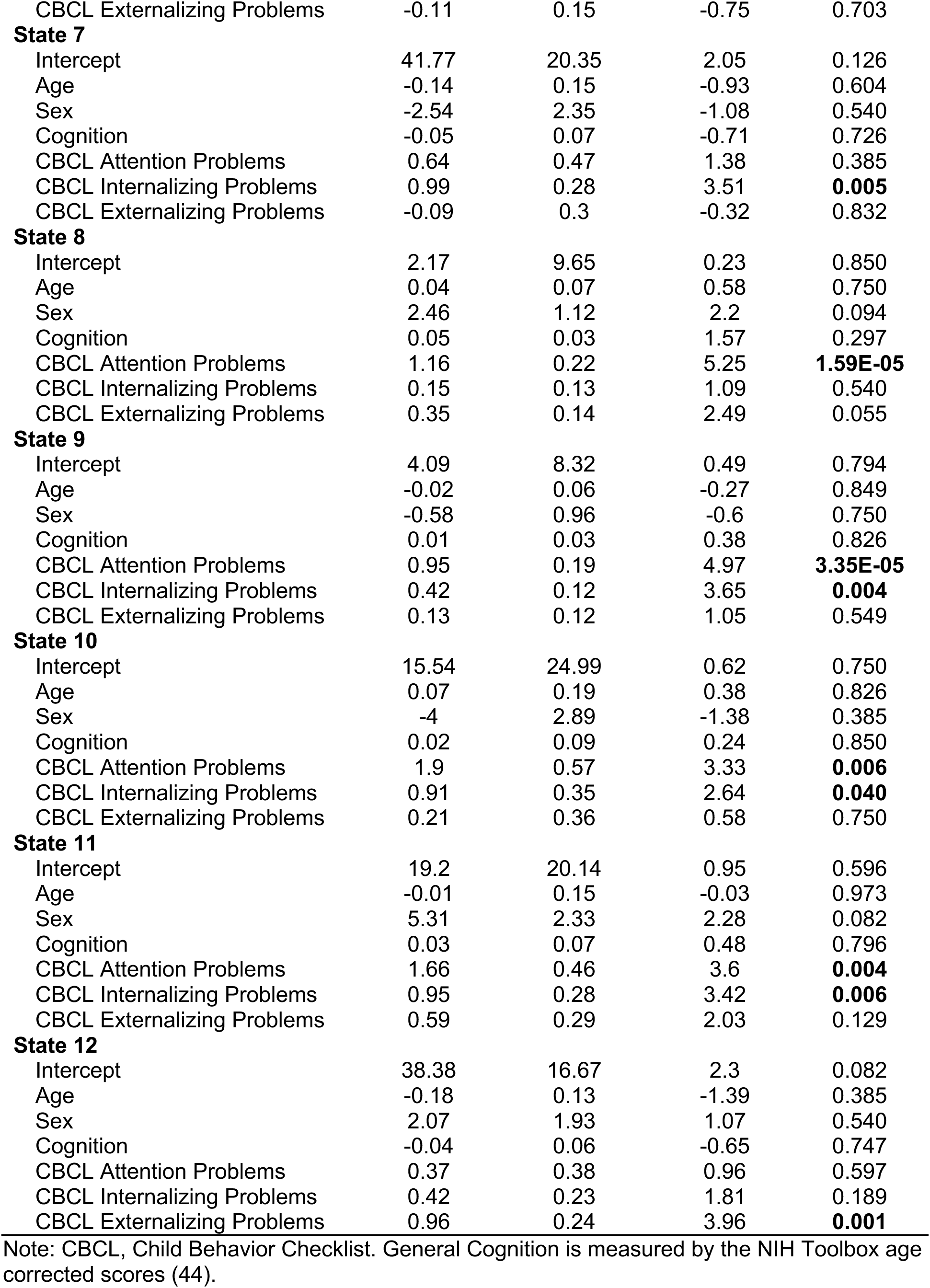
Linear regression models depicting associations between occupancy time and disruptive behavior severity across all latent states.

### Relationship Between Brain State Connectivity and Disruptive Behavior Problem Severity

We then applied multiple regression models to probe the association between state occupancy time (for each latent state) and disruptive behavior problem severity while accounting for commonly co-occurring transdiagnostic symptom domains of attention and internalizing problems. Regression findings across all models are provided in **Table 2**. We found that greater occupancy time in state 12 was uniquely and significantly associated with disruptive behavior severity (*p_FDR_* = 0.001) (**Table 2** and **Figure 3D-E**). For state 12 occupancy time, which was uniquely associated with disruptive behavior problems, a pattern was observed within and between network connectivity among frontoparietal, default mode, dorsal and ventral attention, temporal parietal, sensorimotor, and limbic networks (**Figure 3**). We characterize this pattern in later sections. A highly similar pattern of findings was observed when repeating the primary analyses and accounting for mean head motion (using mean framewise displacement), in which an association emerged between occupancy time for state 12 and disruptive behavior problem severity (**Supplementary Results Table S1**). While our a priori hypotheses focused on network dynamics unique to disruptive behavior problems after accounting for commonly occurring symptom domains, there were also distinct significant associations that emerged for attention and internalizing problems. For the interested reader, we provide a summary of findings for other transdiagnostic domains in the **Supplemental Results** and **Figure S1**.

We then conducted supplemental analyses of state differences to further characterize the connectivity pattern of state 12, which was associated with disruptive behavior severity, in comparison to all other brain states. **Figure 4** shows connectivity differences between state 12 and all other states. Among positive connections, brain state 12 showed weaker connectivity compared to states 1, 4, 6, 7, 9, and 10 across a distributed pattern of networks (**Figure 4A)**. Among negative connections, brain state 12 showed patterns of both stronger connectivity (compared to states 1, 4, 6, 7, 9, and 10) and weaker connectivity (compared to states 2, 3, 5, 8, and 11) primarily in the limbic network (**Figure 4B**).

**Figure 4.**
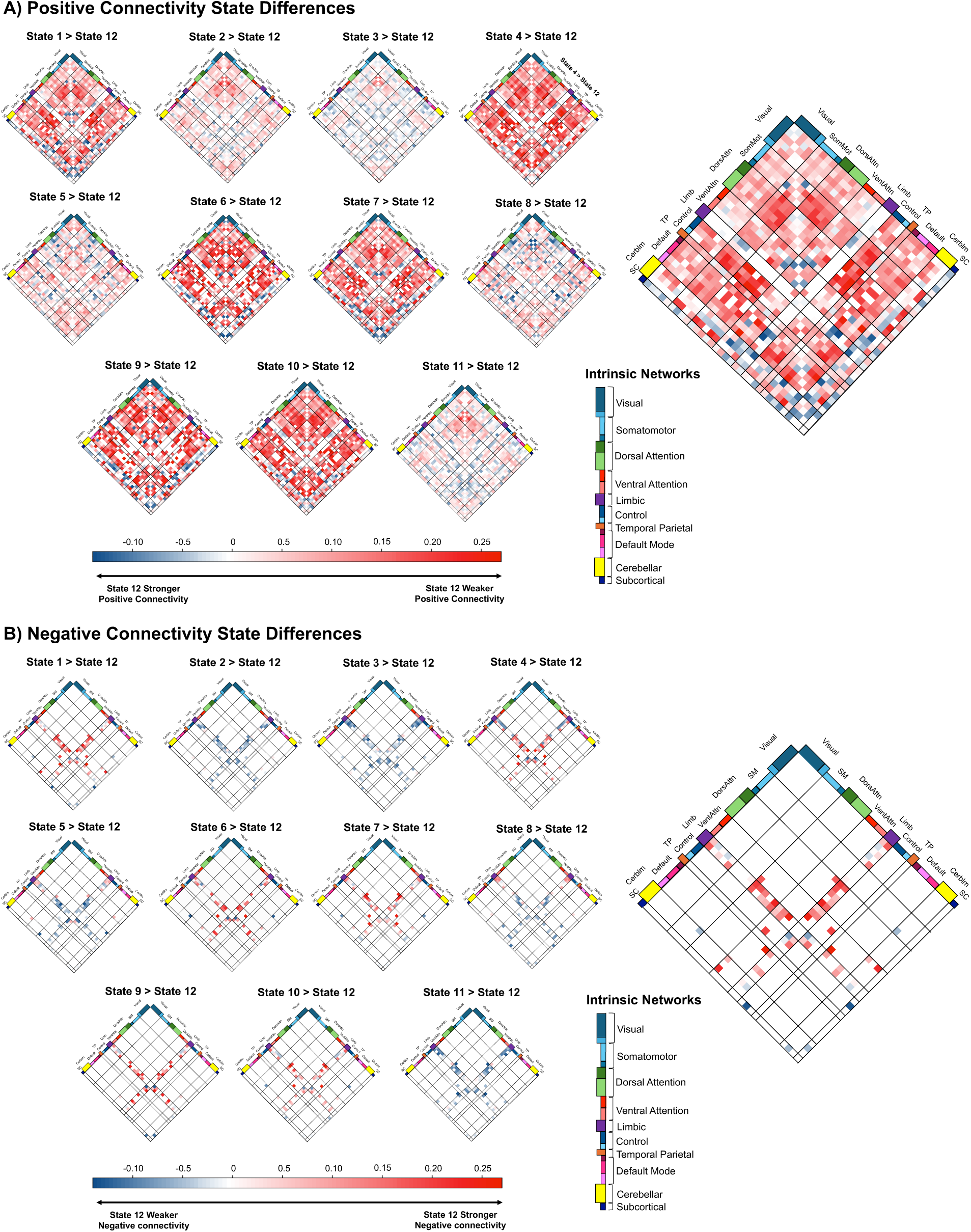
Difference matrices of latent brain states compared to state 12. Given that primary analyses showed a link between occupancy time in state 12 and disruptive behavior severity, additional analyses compared connectivity of state 12 with every other state. Heat maps present the difference in correlation values for each region pair between state 12 vs states 1 through 11. Panel A shows correlation differences for the subset of ROI pairs with a positive correlation within state 12 and the state being compared to state 12. Red cells indicate a stronger positive correlation compared to state 12. Blue cells indicate a weaker positive correlation as compared to state 12. White cells indicate ROI pairs with no difference between the two states. Panel B shows correlation differences for the subset of ROI pairs with a negative correlation in state 12 and the state being compared to state 12. Red cells indicate a weaker negative correlation compared to state 12. Blue cells indicate a stronger negative correlation compared to state 12. White cells indicate ROI pairs with no difference between the two states. Abbreviations for intrinsic networks: Cerblm, cerebellar; Control, Frontoparietal control network; Default, default mode network; DorsAttn, Dorsal Attention Network; Limb, limbic; Visual, visual network; SomMat, somatomotor; SC, subcortical; TP, temporo parietal; VentAttn, Ventral Attention Network.

We also tested for associations between network state dwell or sojourn times (representing time in a given state prior to transitioning to another state) and disruptive behavior problems while accounting for other transdiagnostic symptom domains (attention and internalizing problems). There were no significant associations after FDR-correction between minimum or maximum dwell times for each state and disruptive behavior problems (all *p*s > 0.2) (**Supplemental Results Tables S2-S3** and **Figure S2**). However, for the interested reader, we provide additional details on findings related to attention and internalizing symptoms that were associated with maximum sojourn times in the **Supplemental Results** and **Table S5**.

### Topology of Brain States

To visualize the topography for each of the 12 states, we selected the top 20% of positive correlations while accounting for connections across all states (**Figure 5**). To facilitate interpretation, we further characterized the brain states by estimating metrics of nodal and global topology (**Tables S4**). States 2, 8, and 12 showed high levels of betweenness, while states 1, 4, 6-7, and 9-10 showed low levels. State 12 showed a low level of centrality, while states 6 and 9 showed a high level. State 12 showed low levels of local and global efficiency, while states 9 and 10 showed high levels. State 12 also showed a low level of clustering coefficient, while states 6, 9 and 10 showed a high level. For the interested reader, we also provide graph theory metrics across networks in each state (**Supplemental Tables S5**).

**Figure 5.**
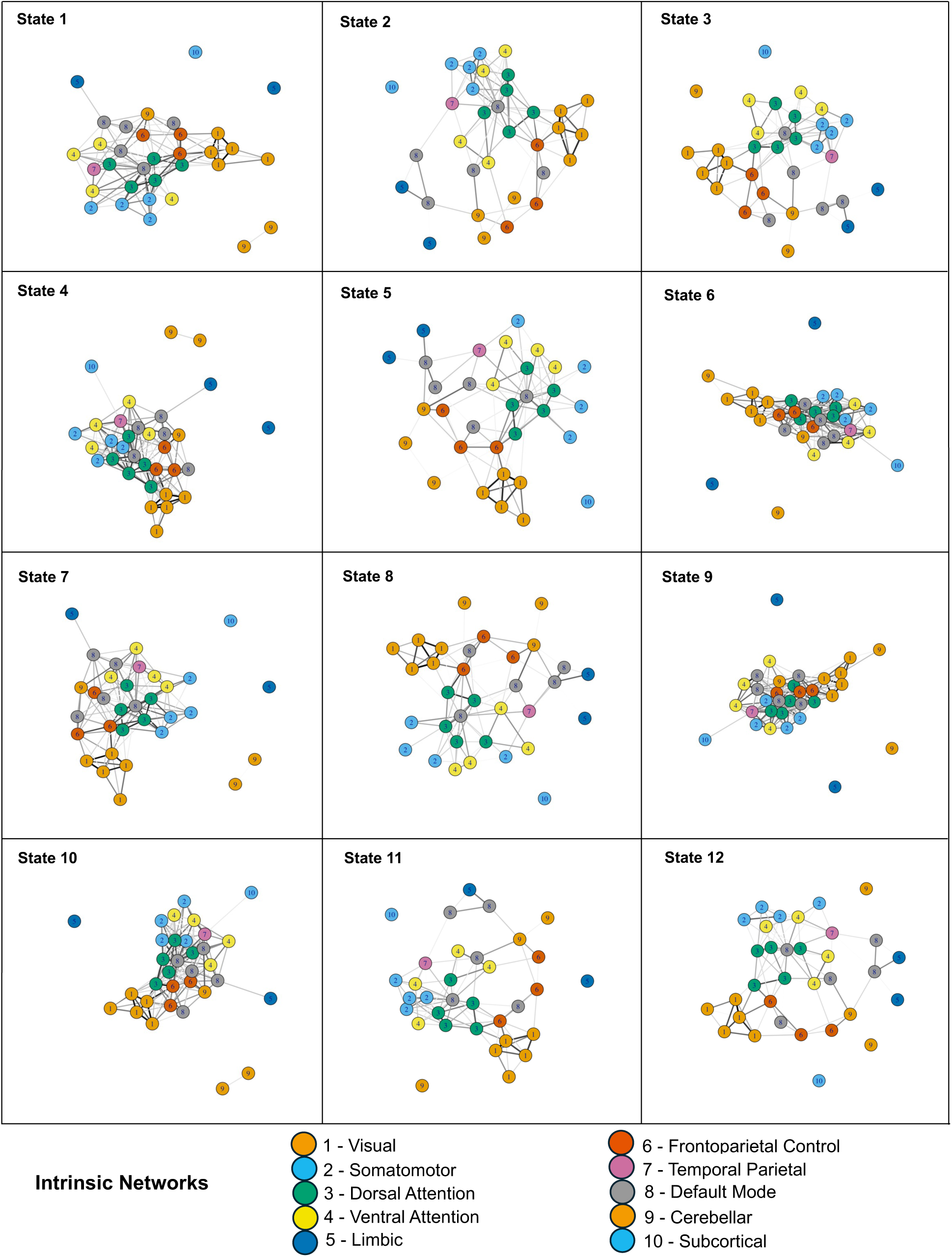
Topological representations of the top quartile of edges for all brain states. For each plot, edge widths are proportional to correlation value (i.e., darker and wider edges indicate greater positive correlations). Vertex color is specific to each intrinsic network.

### Supplemental Follow-Up for Modularity of Networks for Brain State 12

As a follow-up supplemental analysis to help characterize state 12, we examined modularity across independent components and large-scale networks for state 12, which showed a significant, positive association with disruptive behavior problems (**Figure 6**). To facilitate comparison, we also compared state 12 modularity to a state showing an opposite pattern of association with disruptive behavior problems. Thus, we selected state 2 that indicated a negative association with disruptive behavior severity. State 12 consisted of four modules (**Figure 6A-B**), while state 2 consisted of five modules (**Figure 6C-D**). The primary component of the fifth module in state 2 was IC 24, associated with the dorsal attention network. In state 12, the dorsal attention network (IC 24) showed a truncated pattern of connections to modules, while in state 2 there was a more distributed pattern of connections between the dorsal attention network and all other modules (**Figure 6C-D**). Specifically, state 2 showed a stronger connection between the dorsal attention network (IC 24) and the somatomotor network (IC 28, IC 29, and IC 37) compared to state 12. Additionally, state 2 showed stronger connections among somatomotor nodes, including between IC 29 and IC 37 as well as between IC 29 and IC 28.

**Figure 6.**
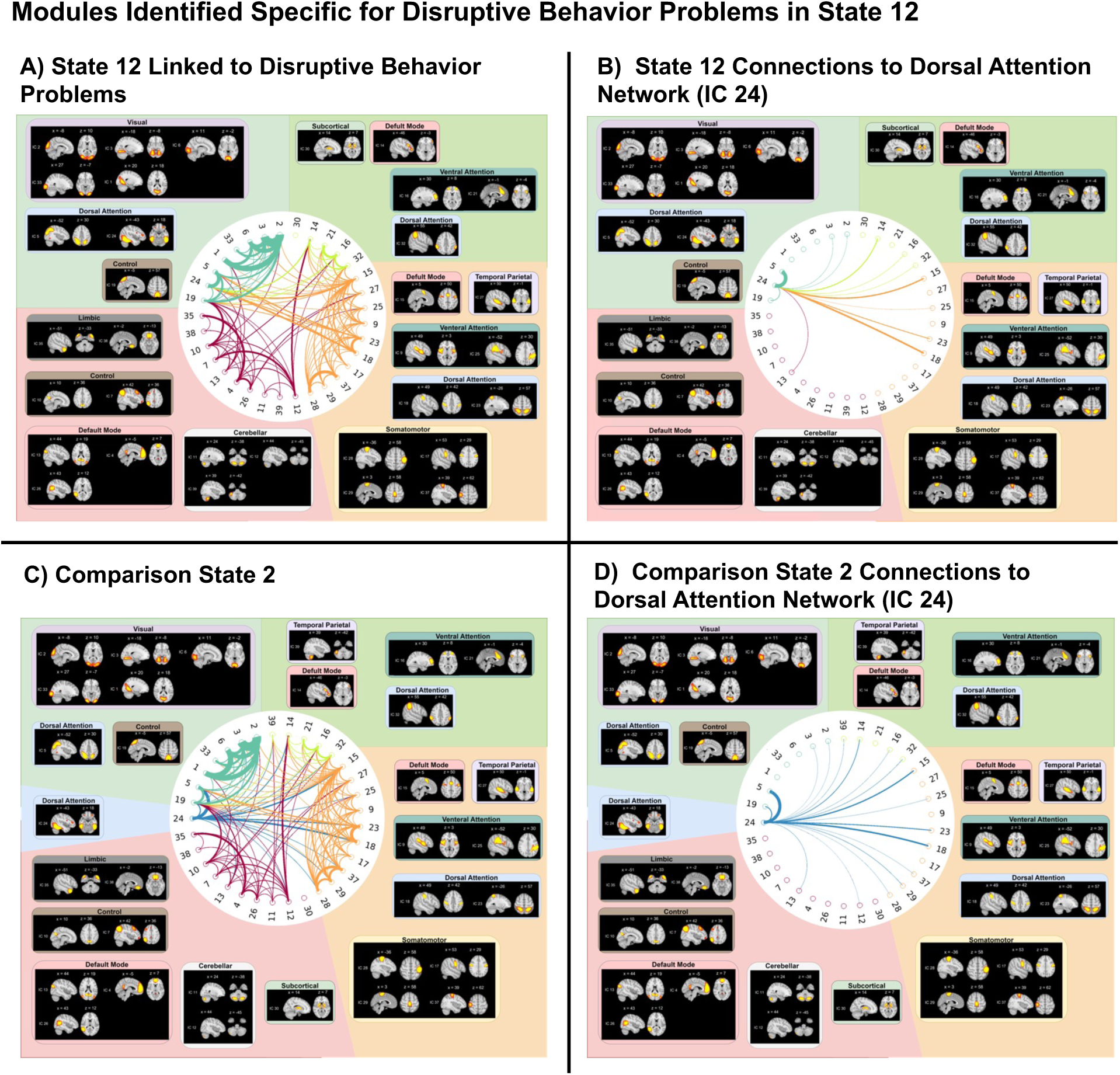
Follow-up supplemental analysis of modularity for states specific to disruptive behavior problems. In primary analyses, for any states emerging that showed significant and distinct or specific associations with disruptive behavior problems, we examined modularity to further understand state characteristics. Thus, we used a modularity analysis to identify community structures within the state. State 12 (A) was significantly and distinctly associated with disruptive behavior problems. To facilitate comparison, we selected state 2 because this state showed an opposite pattern of association with disruptive behavior problems. We conducted modularity analyses on state 12 (primary) (A-B) and state 2 (comparison) (C-D). The dorsal attention network independent component (IC) 24 was selected to spotlight connections with other modules (B and D) for comparison to primary state 12. That is, IC 24 emerged as a unique module in state 2 and showed distinct patterns of connections in state 12 vs state 2.

### Replication of Brain States and Associations with Disruptive Behavior Problems Severity

Thirty-three independent components were retained and 17 were identified as artifacts (**Supplemental Results Figure S5**). Estimated transition probability matrices obtained from fitting the replication HSMM models showed a pattern of attractor states (32) in which states 6-7 and 11 were more likely to be transitioned to from other states (**Figure 7C**). We focused on state occupancy time for replication analyses because this dynamic connectivity metric showed significant and distinct associations with disruptive behavior problems in the discovery dataset. Regression findings across all models are provided in the **Supplemental Results Table S6**. The replication dataset revealed a highly similar pattern of results indicating a positive association between occupancy time and disruptive behavior problem severity. We found that greater occupancy time in states 5 and 6 was uniquely and significantly positively associated with greater disruptive behavior problem severity (**Table 2** and **Figure 7D**). It is important to note that states 5 and 6 are not the same states 5 and 6 from scans 1-2, as a unique set of states was fit to these scans. Therefore, we investigated whether states 5 and 6 from scans 3-4 showed similar characteristics to state 12 from scans 1-2. Additional details regarding regression models and findings for other transdiagnostic domains are provided in the **Supplemental Results Table S6** and **Figure S6**. States 5 and 6 showed a similar pattern of overall weaker connectivity compared to other states (**Supplemental Results Figure S7A-B**), as well as similar topology (**Supplemental Results Figure S7C**) as the discovery analysis. To further characterize states distinctly predicting disruptive behavior problems, we then conducted supplemental analyses of connectivity differences and modularity for brain states 5 and 6. Similar to the primary findings, states 5 and 6 showed weaker connectivity across distributed networks and a truncated pattern of connections between the dorsal attention network (IC 4) and other modules including cerebellar, visual, limbic, default mode, temporoparietal, and frontoparietal control networks (see **Supplemental Results** and **Figure S8**).

**Figure 7.**
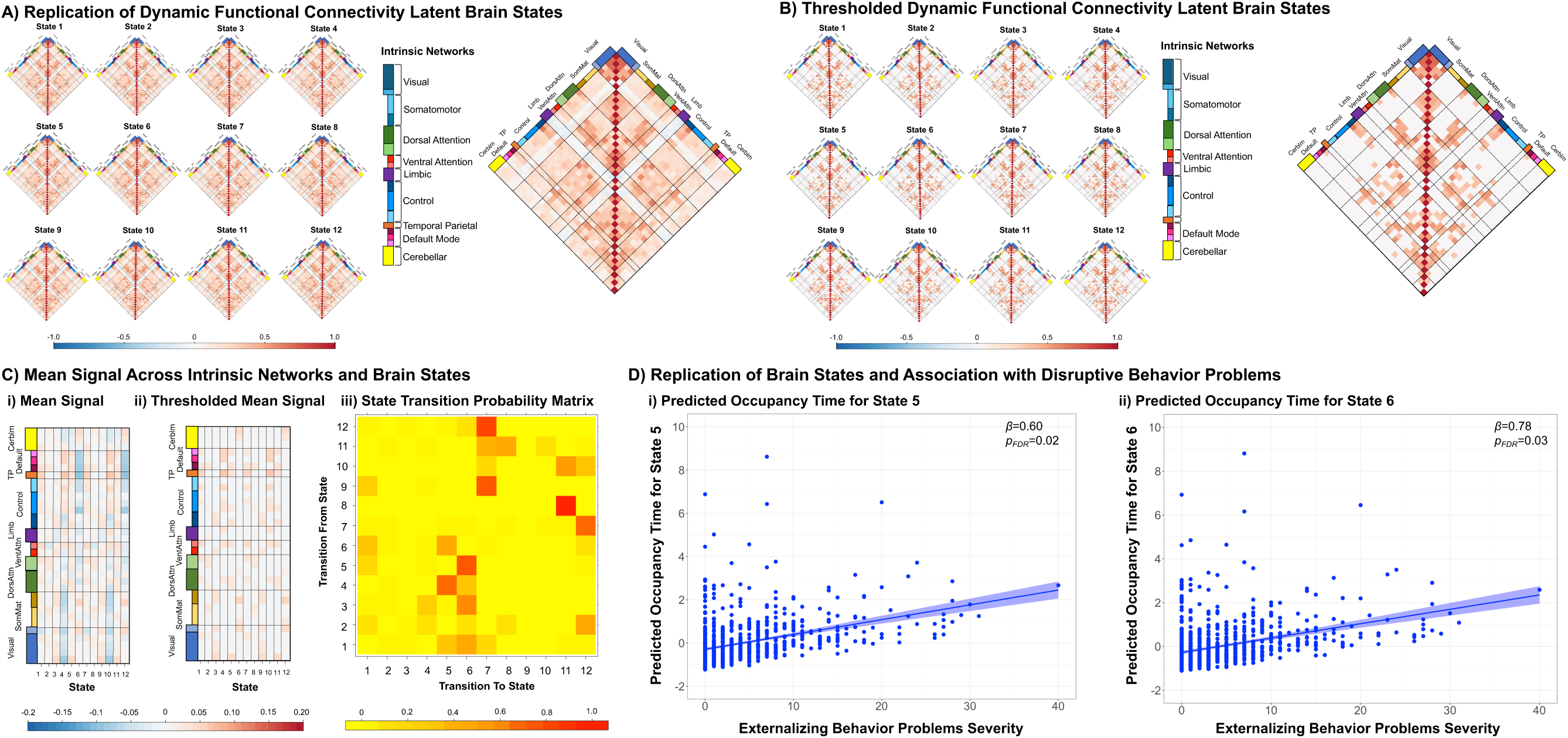
Replication of findings indicates occupancy time in brain states 5 and 6 is associated with severity of disruptive behavior problems. In a replication sample of two held-out resting-state runs per participant, primary analyses were repeated. Mean connectivity is shown for each replication state (A) and after thresholding based on the top quartile of significant edges (B). Red cells indicate a positive correlation while blue cells indicate a negative correlation (or anticorrelation) between two corresponding ROIs. Transition matrices are also shown for replication states, which indicates probabilities that are conditional on switching to a new state (C). Occupancy time in state 5 (Di) and state 6 (Dii) is distinctly positively associated with severity of disruptive behavior problems, which is shown in the regression plots. P values are FDR-corrected across all tests. See Supplemental Results Table S6 for regression model parameters.

## Discussion

This study examined dynamic network connectivity signatures linked to disruptive behavior problems in a transdiagnostic sample of children. Three key findings emerged. First, during resting-state fMRI, greater severity of disruptive behavior problems was uniquely associated with greater time spent (occupancy time) in brain states characterized by overall dysconnectivity, particularly hypo-connectivity. Brain state configurations were identified for 12 states, which mapped onto canonical network parcellations, that also fluctuated around global attractor states. Second, replication of findings was observed in a held-out sample of resting-state runs. Third, occupancy time within large-scale networks implicated in cognitive control (frontoparietal), social perception (default mode), emotion generation (limbic), and attention (dorsal and ventral attention) characterized by globally aberrant connectivity patterns was related to disruptive behavior problems. For the first time, this study demonstrates proof-of-concept for HSMM-derived brain states linked to severity of disruptive behavior problems in children. That is, we identify a potential expression of core canonical, time-varying network configurations or brain states that may be uniquely linked to disruptive behavior problems in children. It is possible that resulting aberrant, globally disconnected brain states may disrupt cognitive control circuitry supporting related psychological processes of executive functioning, emotion regulation, and inhibitory control—leading to increased severity of disruptive behavior problems. These findings also lend support to a broader network dysfunction model of childhood disruptive behavior problems, which posits that perturbations across cognitive control, social perception, emotion generation, and sensorimotor networks are implicated in the risk and onset of maladaptive aggression (10).

### Disruptive Behavior Problems and Cognitive Control Network Dynamics

In the current study, children with elevated levels of disruptive behavior problems showed state-specific, intermittent disruptions within cognitive control networks supporting the top-down control of emotion as well as broader psychological processes including inhibitory control and executive functioning. We applied HSMM to identify dissociable network configurations in 12 brain states distinctly linked to transdiagnostic disruptive behavior problems in youth. Of note, we observe that dynamic functional connectivity, particularly related to occupancy time (i.e., the time spent in a state), is unique or symptom-specific and relatively stable or reliable in predicting disruptive behavior severity over the course of a scanning session. That is, distinct brain states continued to show unique relationships with disruptive behavior problems across held-out resting-state runs within the scanning session. In support of this, patterns of dynamic connectivity are heritable (35, 64) and replicable (32, 65)—similar to networks defined through traditional static analyses—and show relationships with cognitive traits as well as trait-like signatures of psychopathology (26, 31, 32, 65–68). Recent work suggests that temporal fluctuations in network connectivity across discrete brain states may also be anchored in neuroanatomic connectivity (69, 70) with time-varying, transient configurations of functional networks reflecting the impact of psychological or cognitive processes (19, 21). Our findings are consistent with prior evidence suggesting alterations in dynamic brain architecture linked to globally disconnected states in clinical populations of children and adolescents, which could suggest that disruptions across association networks, particularly cognitive control circuitry, in child mental health may emerge via perturbations in transient network configurations (25, 66, 68, 71–73).

Given the emerging evidence linking symptom domains to temporally dynamic configurations of large-scale networks (24–26, 30, 66–68, 71–76), the presence of potential ‘fingerprints’ derived from brain dynamics linked to network-specific impairment and psychopathology has implications for developing brain-based biomarkers to inform clinical practice. Recent work suggests promising findings in the implementation of dynamic neural markers for assessing treatment outcomes in depressive and schizophrenia spectrum disorders (26, 65, 77–79). For instance, dynamic neural markers could be combined with structural and/or static connectivity neural markers to provide complementary data for prediction of brain-behavior associations, provide a potential individual-specific marker for identifying symptom dimensions to assist with diagnosis and early identification, and/or used to assess target engagement and response to treatment, which could inform treatment decisions and selection of individually-specific intervention modalities (e.g., cognitive-behavioral intervention, psychotropic medication, or a combination of both).

Children with greater severity of disruptive behavior problems spent greater time in states characterized by hypo-connectivity and reduced modularity (i.e., less segregation of sub-networks). Additionally, follow-up modularity analyses indicated that state 12 showed weaker connections between the dorsal attention network and other networks. Altered activity and connectivity within the dorsal attention network is implicated in disruptive behavior problems (15, 80, 81). Further, attention, frontoparietal, and default mode networks are among the most globally connected in the brain and characterized as core hubs of integration for cognitive control and social perception processes (82, 83). Here, findings suggesting a less distributed pattern of connections among the dorsal attention network in state 12 linked to disruptive behavior severity could suggest reduced segregation of cognitive control sub-networks, which aligns with prior findings of reduced modularity and segregation of networks in ADHD (31), schizophrenia (76), and ASD (73).

Findings of increased occupancy time associated with severity of disruptive behavior aligns with prior work reporting reduced variability in dynamic networks, including less frequent transitions between states and/or longer dwell times in globally disconnected states, linked to child psychopathology including ASD (25, 75, 84–88), ADHD (31, 66, 68), and depression (71). Decreased transitions may indicate overly stable dynamic properties linked to cognitive inflexibility, which could impede shifting of psychological processes critical for cognitive control. In support of this, findings from prior studies have suggested increasing variability (i.e., increase in state transitions) across age (89, 90). Furthermore, decreasing dwell time is thought to be associated with increasing age across the lifespan (89, 90) whereby greater dynamic variability with age (marked by frequent transitions across states) may reflect greater complexity and enhanced capacity for cognitive control processes including inhibitory control, information processing, cognitive flexibility/shifting, and executive functioning (91). That is, increased dynamic variability with age could afford increased levels of neurocognitive and behavioral flexibility. Thus, our finding of increased occupancy time in states associated with disruptive behavior severity could suggest perturbations in cognitive control, which may potentially hinder the successful top-down regulation of emotion and impede cognitive flexibility (e.g., in response to frustration-inducing situations). Alternatively, increased dwell time in globally disconnected states linked to increased symptom severity could indicate attenuated maturation of networks and/or reduced integration across networks supporting cognitive control such as frontoparietal and attention circuitry (30). Future longitudinal studies will be necessary to understand the trajectories of dynamic networks and how this corresponds to development and risk of onset for child psychopathology.

### Limitations

First, the current study was cross-sectional. Future studies leveraging the ABCD Study as well as clinical samples will be important to understand how longitudinal trajectories and disruptions in the reconfigurations of dynamic networks across development are linked to emergence and risk of onset for disruptive behavior problems in youth. Second, findings reported here are related to resting-state fMRI and may not generalize to dynamic connectivity associations during task-based fMRI. Therefore, additional studies leveraging task-based dynamic functional connectivity models will help to assess non-task versus task reconfigurations and associations with symptom severities related to state versus trait features (92). Third, dynamic network analyses of fMRI data are limited by the relatively slow hemodynamic BOLD response. Comprehensive understanding related to time-varying properties and transient changes occurring on faster timescales would benefit from integration with multimodal approaches including electrophysiological methods (e.g., EEG, MEG), which would complement fMRI in terms of spatial resolution and brain coverage. Lastly, the sample size for this dataset is modest, limiting statistical power. However, we selected a modest sample as a proof-of-concept with considerations for computational load. Future studies leveraging computationally ‘inexpensive’ approaches will be important for developing efficient and generalizable analytical workflows to derive dynamic networks. Nonetheless, the present study constitutes an initial step in characterizing time-varying, dynamic patterns of connectivity in youth that are linked to transdiagnostic symptom domains.

## Conclusion

The functional chronnectome has potential to enhance understanding and identification of brain-based biomarkers to characterize and typify the underlying circuit mechanisms of transdiagnostic symptom domains in youth, particularly disruptive behavior problems. These findings demonstrate that disruptions in the time-varying properties of functional connectivity are associated with disruptive behavior problems and could serve as a potential neural marker of individual differences in DBD. Dynamic functional connectivity approaches may provide vital information to help predict transdiagnostic symptom dimensions in youth that could accelerate identification of robust brain-based biomarkers to inform individualized treatments and clinical decisions.

## Acknowledgments

K.I. is supported by the National Institute of Mental Health (K23-MH128451). This work was supported by National Center for Advancing Translational Sciences grant KL2 TR001862 (K.I.) and TL1 TR001864 (K.I.), a Yale Child Study Center Junior Faculty Development Pilot Award (K.I.), and the Yale Child Study Center Translational Developmental Neuroscience Training Program (T32 MH18268) (K.I.). H.M.S is supported by the National Institute of Biomedical Imaging and Bioengineering (K25-EB032903-01).

## ABCD acknowledgement

A portion of the data used in the preparation of this article were obtained from the Adolescent Brain Cognitive Development (ABCD) Study (https://abcdstudy.org), held in the NIMH Data Archive (NDA). This is a multisite, longitudinal study designed to recruit more than 10,000 children ages 9-10 and follow them over 10 years into early adulthood. The ABCD Study® is supported by the National Institutes of Health and additional federal partners under award numbers U01DA041022, U01DA041028, U01DA041048, U01DA041089, U01DA041106, U01DA041117, U01DA041120, U01DA041134, U01DA041148, U01DA041156, U01DA041174, U24DA041123, U24DA041147, U01DA041093, and U01DA041025. A full list of supporters is available at https://abcdstudy.org/federal-partners.html. A listing of participating sites and a complete listing of the study investigators can be found at https://abcdstudy.org/consortium_members/. ABCD consortium investigators designed and implemented the study and/or provided data but did not necessarily participate in analysis or writing of this report. This manuscript reflects the views of the authors and may not reflect the opinions or views of the NIH or ABCD consortium investigators. The ABCD data repository grows and changes over time. The ABCD data used in this report came from NDA 4.0 release (DOI: <u>10.15154/z563-zd24</u>). We are grateful to the study participants for their time and participation.

## Preprint Servers

A version of this manuscript was posted as a preprint on bioRxiv. The authors retain full copyright.

