## Supplemental Information for "Dynamic Resting-State Network Markers of Disruptive Behavior Problems in Youth"

^1^Wake Forest University School of Medicine, Department of Biostatistics and Data Science, ^2^Yale University School of Medicine, Child Study Center, ^3^Yale University School of Public Health, Department of Social and Behavioral Sciences, ^4^Virginia Tech-Wake Forest University School of Biomedical Engineering and Sciences, ^5^Yale University, Department of Psychology, ^6^Yale University, Wu Tsai Institute, ^7^Johns Hopkins University, Bloomberg School of Public Health

***Please address correspondence to**

Karim Ibrahim, Psy.D.

Assistant Professor in the Child Study Center and of Psychology

Yale University School of Medicine

Child Study Center

230 South Frontage Road

New Haven, CT 06520

**Supplemental Background:**

Recent advances in functional connectivity indicate that large-scale networks exhibit time-varying properties that can be linked to brain-behavior associations, which has potential to provide new insights into the underlying circuit mechanisms of psychopathology (1-3). Thus, there are several advantages to dynamic connectivity approaches, which complement information gained from static connectivity approaches. First, measures of dynamic connectivity reveal the moment-to-moment reconfigurations of networks without implicit assumptions that connectivity is static (1, 4, 5). Second, time-varying measures of connectivity could represent a complementary approach for characterizing the dynamic nature of the connectome. For instance, human cognition is a fluctuating process and connectivity patterns exhibit complex spatiotemporal dynamics at multiple time scales (6). Third, while static connectivity approaches have played a central role in elucidating the circuitry underlying disruptive behavior (7), dynamic connectivity also holds promise in advancing development of neural markers of psychopathology (8, 9) in diverse pediatric clinical populations (10), particularly related to childhood disruptive behaviors.

**Supplemental Methods:**

**Denoising of Imaging Data**

Data were corrected for structured noise associated with motion artifacts using the AROMA package (11) (https://github.com/rhr-pruim/ICA-AROMA). AROMA is an independent component analysis (ICA) based method that automatically classifies and removes components identified as noise. ICA-AROMA is a robust approach for denoising and removing motion artifact in pediatric fMRI data (12, 13) and preserving temporal degrees of freedom while eliminating distance-dependent artifact without inducing false anti-correlations (11, 12). Noise components were detected and removed prior to any temporal filtering so the resulting cleaned data were temporally high-pass filtered. At the single-subject level, we also regressed the white-matter timeseries, which was included as a confound variable in analyses. Denoised resting-state fMRI data or timeseries were used for consequent preprocessing and analysis steps described in the main text.

**FreeSurfer Preprocessing and Analysis**

Standard preprocessing and analysis was conducted using FreeSurfer v6.0 (http://surfer.nmr.mgh.harvard.edu) surface-based cortical reconstruction (14, 15). This involved cortical surface reconstruction, cortical thickness estimation, cortical parcellation, and subcortical segmentation. The surface-based morphometry approach employs information related to intensity and continuity from the entire 3D MRI volume in segmentation and deformation procedures (16). This approach also uses spatial intensity gradients across tissue classes (vs reliance upon absolute signal intensity) (15). The surface reconstruction and estimation of metrics related to the white matter surface (comprising the white-gray matter interface) and the pial surface (comprising the gray matter-cerebrospinal fluid interface) have been described and validated in prior studies (16-18).

**Imaging Processing**

Minimally preprocessed resting-state fMRI data were processed using Functional MRI of the Brain (FMRIB) Software Library (FSL Version 4.1.6; FMRIB, Oxford, United Kingdom) (19, 20) and denoised using the AROMA package (11) (<https://github.com/rhr-pruim/ICA-AROMA>). The first six volumes of each functional run were discarded to allow for the magnetization to reach a steady state. Motion was corrected using FSL MCFLIRT linear realignment tool (21). Functional data were temporally realigned to correct for interleaved slice acquisition. Data were spatially smoothed with a 5-mm full width at half maximum isotropic Gaussian kernel with a non-linear high-pass filter (100 s cut-off). Individual participant analyses were conducted using FSL FMRI Expert Analysis Tool (FEAT). Functional images were registered to high-resolution T1-weighted images and normalized to the Montreal Neurological Institute 152 template through an affine transformation and a nonlinear warp.

**Independent Component Analysis**

Large-scale intrinsic networks were identified at the group level based on a data-driven approach using FSL's Multivariate Exploratory Linear Optimized Decomposition into Independent Components (22) (MELODIC) tool (v3.15). A multisession temporal concatenation approach was used to generate a parcellation of 50 spatiotemporal components covering both the cortex and subcortex. To ensure robustness of intrinsic networks, independent components were first visually inspected by three experienced neuroimagers (K.I., G.H., G.M.). Consensus was reached for each component. Next, ICA components were mapped to Yeo intrinsic networks as in previous work (23). We statistically compared the spatial map of each ICA component to a set of 17 networks based on the Yeo parcellation (24) (visual networks A-B, somatomotor networks A-B, temporal parietal network, dorsal attention networks A-B, salience networks A-B, executive control/frontoparietal networks A-C, default mode networks A-C, limbic networks A-B). We used FSL’s “fslcc” tool to calculate Pearson's *r* for each pairwise relationship and kept only those ICA components that yielded a significant spatial correlation (*r* > 0.3) with one of the networks from Yeo et al. (24). We also implemented maps for cerebrospinal fluid, corpus callosum, and white matter, and ICA components correlating with these features were excluded. Subcortical regions were assigned to subcortical network (e.g., amygdala, hippocampus, thalamus). Cerebellar regions were assigned to a cerebellar network. Thus, 17 ICA components were determined to be motion or artifact (e.g., white matter, cerebrospinal fluid), leaving 33 ICA components of high quality included in the final analysis. Time series were then extracted for the 33 cortical and subcortical ICA components.

**ABCD Study inclusion variables/flags**

The inclusion criteria for each imaging modality are aligned with the “ABCD Release 4.0 release notes” available at DOI 10.15154/1523041: “NDA 4.0 MRI Quality Control Recommended Inclusion”. For more detail regarding the ABCD Study structural and resting-state sequences and acquisition parameters, see Casey et al. (25) and Hagler et al. (26). Based on ABCD imaging analysis recommendations and DAIRC inclusion flag (“imgincl_rsfmri_include” of the “abcd_imgincl01”), participants in the current study passed protocol compliance and quality control for T1-weighted and resting-state scans. We provide a summary from the release notes of these inclusion criteria related to resting-state as follows:

**T1-weighted (variable *imgincl_t1w_include* = 1).** Must pass both raw QC for T1 and FreeSurfer QC. Derived results must exist.

**Resting-state fMRI (variable *imgincl_rsfmri_include* = 1).** Must pass raw QC and T1 QC, with more than 375 frames post-censoring. B0 unwarp must be available. FreeSurfer QC and manual post-processing QC must be successful. Registration to T1w must be less than 19. Dorsal cutoff score must be less than 65, and ventral cutoff score less than 60. Derived results must exist.

**ABCD Study QC criteria**

The passing QC criteria for each imaging modality are summarized here from the “ABCD Release 4.0 release notes” available at DOI 10.15154/1523041: “NDA 4.0 MRI Quality Control Recommended Inclusion”. For more detail regarding the ABCD Study QC criteria, scan sequences, and/or imaging acquisition parameters, see Casey et al. (25) and Hagler et al. (26). We provide a summary from the release notes of these QC criteria related to resting-state as follows:

**T1-weighted.** Passing raw QC is assessed based on the following signs of artifacts and poor imaging quality: presence of wrap-around field of view artifacts, brain cut-off due to the participant motion outside prescribed slices, magnetic susceptibility artifacts due to dental implants and T1w and T2w motion artifacts (e.g., blurring and ghosting). FreeSurfer QC is assessed based on the severity of five types of artifacts or processing problems: motion, intensity inhomogeneity, white matter underestimation, pial overestimation and magnetic susceptibility artifact. Values are assigned ranging from 0 to 3 (absent, mild, moderate or severe artifact respectively). An overall QC score of 1 indicates a pass based on all items having a score between 0-2.

**fMRI.** Passing raw and FreeSurfer QC as detailed above. Functional images are assessed based on the severity of five types of artifact or processing problems: B0 warping, magnetic susceptibility artifact, full head coverage, and T1w registration. Values are assigned ranging from 0 to 3 (absent, mild, moderate, or severe artifact, respectively). An overall QC score of 1 indicates a pass based on all items having a score between 0-2.

**Topological Characteristics of Brain States**

Exploratorily, we implemented graph theory to derive metrics of degree centrality, betweenness, local efficiency, and clustering coefficients for each state and networks comprising states. Global and nodal properties were estimated for each state and networks: Visual (VIS), Somatomotor (SM), Dorsal Attention Network (DAN), Ventral Attention Network (VAN), Limbic (LIMB), Frontoparietal control (CON), Default Mode Network (DMN), Cerebellum (CEREBLM), and Temporoparietal (TP). We present these values in the **Supplemental Results**.

**HSMM Complete Data Log-Likelihood**

Our previously validated HSMM framework for inferring time-varying brain networks from fMRI data estimates a set of network states, dwell/sojourn time distributions, and probabilities for switching from one state to another – all at the group level (10, 27). Individual estimates of state sequences can be further derived by running a Viterbi algorithm (58), and those state sequences can be used to obtain individual estimates of overall time spent in each state, as well as transition probabilities. Briefly, we denote the ICA time-series data for each participant by $Y_{i1}, \ldots, Y_{iT}$, where each $p$−dimensional vector $Y_{iT}$ contains the BOLD measurements of the $p$ICs at the $t^{th}$ timepoint for the $i^{th}$ participant. The collection of vectors of the observed time-series data is denoted by $\tilde{Y}_{i}$. Now suppose a unique brain network state gives rise to each $Y_{it},$ but we cannot observe this. We represent this hidden network index variable underlying the observed time-series vectors at a particular observation time by $S_{it},$where $S_{it}$ takes on discrete values. That is, $S_{it}, \in\left\{ 1,\ldots, K \right\}.$The vector of hidden network state variables, $S_{i1}, \ldots, S_{iT}$, for a single individual is denoted by $\tilde{S}_{i}$ . We assume that each $Y_{it}$follows a multivariate Gaussian distribution $Y_{it} \sim N\left( \mu_{s}, \Sigma_{s} \right),$ where the mean and covariance depend on the current (unknown/hidden) network state. Thus, each network state has unique mean activations across ICs, and its own covariance structure between ICs. The complete data log-likelihood of the HSMM for one participant, assuming $K$ unique network states, can be written as:

$$\log P(\tilde{Y}_{i}=\tilde{y}_{i},\tilde{S}_{i}=\tilde{s}_{i};\mu_{1:K,}\Sigma_{1:K}, P,d_{1:K})=log f\left( \tilde{s}_{i},\tilde{y}_{i} \right)$$

$=\log f\left( \tilde{y}_{i}|\tilde{s}_{i} \right)+\log f\left( \tilde{s}_{i} \right)$

$=\sum_{t=1}^{T} \log f\left( y_{it} | s_{it} \right)+ \sum_{r=2}^{R} \log(f(s_{ir}|s_{i(r-1)}) d_{s_{ir}}(u_{ir}))$

$+ \log(f(s_{iR}|s_{i(R-1)}) D_{s_{iR}}(u_{iR})) \log(f\left( s_{i1} \right)d_{s_{i1}}(u_{i1}))$,

where $R$ = total number of network state changes + 1, $f\left( s_{1} \right)$ is the probability of starting in a particular state, $d(u)$ is the sojourn time density, $s_{r}$ is the $r^{th}$ visited state, and $u_{r}$is the time spent in that state. The first term is based on the conditional distribution of the observed BOLD signal vector given the underlying hidden network, which takes on a Gaussian distribution, as noted above. The second portion of the equation is made up of two parts. The first is a transition probability matrix, denoted $P$, where the probability at row q and column v represents the probability of transitioning from network state $q$ to $v$ (i.e., $p_{qv}=P(s_{r}=v, s_{r-1}=q$)). The second part, $d_{s_{r}}\left( u_{r} \right),$ represents the dwell time/sojourn distribution. The third portion accounts for the last state a participant enters. Note that

$D\left( u \right)= \sum_{V\geq u} d(v)$

is the survivor function and pertains only to the sojourn time in the final state. It allows us to avoid the assumption that the process is leaving the final state immediately after time $T.$ Lastly, the fourth term accounts for an individual’s initial network state. The log-likelihood is then summed across subjects, and a Maximization Expectation (E-M) algorithm is used to estimate the model parameters.

**Replication Analysis**

To assess the robustness of findings across the sample, we tested replication of findings in a held-out set of two resting-state runs for each subject. That is, the HSMM and linear regression models were repeated using the held-out, two runs of resting-state to identify brain states. Large-scale functional networks were derived using the identical approach above. Thirty-three independent components were retained and 17 were identified as artifact. Twelve network states were estimated using the HSMM, which comprised of multiple large-scale networks with distributed patterns of within and between network connectivity, common to brain-wide connectivity approaches. Identical methods were used to derive 6 states from each subgroup (i.e., 6 states from the low problem behavior group and 6 from the high problem behavior group) to ensure equal representation of states across all behavior scores in the HSMM models. Metrics of dynamic connectivity were implemented in regression models to test replication of brain-behavior associations. We then applied multiple regression models to test the association between dynamic brain metrics for each brain state and disruptive behavior problem severity while controlling for age, sex, motion, cognitive performance and common comorbid transdiagnostic symptoms of attention and internalizing problems.

**Supplemental Results:**

**Exploratory Graph Theory Metrics for Brain States and Networks**

**Table S5** shows values from exploratory graph theory metrics. States 10, 6 and 9 that showed higher degree, and states 5 and 8 showed lower degree centrality. The dorsal attention network showed higher degree across states state 10, 6 and 9. For local efficiency, state 12 showed higher local efficiency, particularly for the dorsal attention network; additionally, states 10, 4, 6, and 9 showed higher local efficiency while state 3 showed lower. Within the default mode network for local efficiency, states 10, 6, and 9 showed higher efficiency and state 3 showed lower. For clustering coefficient, in the dorsal attention network, states 10, 6, and 9 showed higher clustering and state 12 showed lower clustering, including within the default mode network.

**Network Dynamics of Transdiagnostic Symptom Domains of Attention and Internalizing Problems**

While our a priori hypotheses focused on network dynamics unique to disruptive behavior problems after accounting for commonly occurring symptom domains, there were also distinct significant associations that emerged for attention and internalizing problems. Specifically, greater attentional-related problem severity was uniquely associated with reduced occupancy time in states 3 and 6 (*p_FDR_* = 0.02 and *p_FDR_* = 0.03, respectively) (**Table 1** and **Figure S1**), which remained significant after additionally accounting for motion (all *ps* < 0.004) (**Table S1**). Additionally, greater internalizing problem severity was uniquely and significantly associated with reduced occupancy time in state 4 and greater occupancy time in state 7 (*p_FDR_* = 0.006 and *p_FDR_* = 0.005, respectively) (**Table 1** and **Figure S1**), which also remained significant after accounting for motion (all *ps* < 0.002) (**Table S1**). States specific to attention problems showed a pattern of hypo-connectivity (state 3) and hyper-connectivity (state 6) relative to state 12 that was specific to disruptive behavior problems (**Figure 4**). States specific to internalizing problems showed a pattern of hyper-connectivity (states 4 and 7) relative to state 12 that was specific to disruptive behavior problems (**Figure 4**). For the interested reader, we also provide supplemental findings related to network topology of these states (**Figure 5**) and exploratory metrics of graph theory from global and nodal properties (**Supplemental Results** and **Tables S4-S5**).

**Sojourn Time or Dwell Time Associations with Transdiagnostic Symptom Domains of Attention and Internalizing Problems**

While our a priori hypotheses focused on network dynamics unique to disruptive behavior problems after accounting for commonly occurring symptom domains, there were also distinct significant associations that emerged for attention and internalizing problems for sojourn time. For minimum sojourn time, no significant associations emerged for attention or internalizing symptoms (all *p*s > 0.1) (**Table S2** and **Figure S3**). For maximum sojourn time, significant associations emerged for attention and internalizing symptoms (**Table S3** and **Figure S4**). Specifically, greater attentional-related problem severity was uniquely associated with greater sojourn time in state 8 (*p_FDR_* = 0.01) (**Table S3** and **Figure S4**). Additionally, greater internalizing problem severity was uniquely and significantly associated with reduced sojourn time in state 4 (*p_FDR_* = 0.04) and state 5 (*p_FDR_* = 0.01), and greater sojourn time in state 7 (*p_FDR_* < 0.001) and state 9 (*p_FDR_* < 0.001) (**Table S3** and **Figure S4**).

**State Differences for Replication Analysis**

To further characterize states distinctly predicting disruptive behavior problems, we then conducted supplemental analyses of connectivity differences for brain states 5 and 6, in comparison to all other replication brain states. **Figure S7** shows connectivity differences between states 5 and 6, and all other states, which revealed a similar pattern to the discovery analysis. For positive connectivity, brain states 5 and 6 showed a pattern of weaker connectivity across distributed networks compared to all other states (**Figure S7**). For negative connectivity, brain states 5 and 6 showed patterns of weaker connectivity primarily in the limbic network compared to all other states (**Figure S7**).

**Table S1.** Linear regression models depicting associations between occupancy time and disruptive behavior severity across all latent states accounting for motion

| ***Predictors*** | ***β*** | ***Std. Error*** | ***t*** | ***p_FDR_*** |
| --- | --- | --- | --- | --- |
| **State 1** |  |  |  |  |
| Intercept | -0.62 | 14.36 | -0.04 | 0.986 |
| Age | 0.06 | 0.11 | 0.57 | 0.757 |
| Sex | -2.05 | 1.66 | -1.24 | 0.392 |
| Cognition | -0.01 | 0.05 | -0.14 | 0.947 |
| Motion | 29.44 | 2.61 | 11.27 | 2.13E-15 |
| CBCL Attention Problems | -0.71 | 0.33 | -2.17 | 0.074 |
| CBCL Internalizing Problems | -0.03 | 0.20 | -0.16 | 0.942 |
| CBCL Externalizing Problems | -0.11 | 0.21 | -0.51 | 0.757 |
| **State 2** |  |  |  |  |
| Intercept | 97.41 | 26.64 | 3.66 | 0.001 |
| Age | 0.06 | 0.20 | 0.28 | 0.872 |
| Sex | -0.14 | 3.07 | -0.05 | 0.986 |
| Cognition | 0.00 | 0.09 | 0.03 | 0.991 |
| Motion | -60.68 | 4.84 | -12.53 | 2.13E-15 |
| CBCL Attention Problems | -1.70 | 0.61 | -2.80 | **0.016** |
| CBCL Internalizing Problems | -1.38 | 0.37 | -3.76 | **0.001** |
| CBCL Externalizing Problems | -0.85 | 0.39 | -2.20 | 0.070 |
| **State 3** |  |  |  |  |
| Intercept | 96.29 | 21.08 | 4.57 | 3.17E-05 |
| Age | -0.07 | 0.16 | -0.45 | 0.768 |
| Sex | -1.09 | 2.43 | -0.45 | 0.768 |
| Cognition | -0.14 | 0.07 | -1.90 | 0.135 |
| Motion | -46.17 | 3.83 | -12.05 | 2.13E-15 |
| CBCL Attention Problems | -1.20 | 0.48 | -2.49 | **0.037** |
| CBCL Internalizing Problems | -0.66 | 0.29 | -2.27 | 0.062 |
| CBCL Externalizing Problems | -0.52 | 0.31 | -1.71 | 0.185 |
| **State 4** |  |  |  |  |
| Intercept | 61.33 | 19.74 | 3.11 | 0.007 |
| Age | 0.05 | 0.15 | 0.33 | 0.836 |
| Sex | 1.08 | 2.28 | 0.47 | 0.768 |
| Cognition | 0.03 | 0.07 | 0.45 | 0.768 |
| Motion | -23.95 | 3.59 | -6.67 | 4.37E-10 |
| CBCL Attention Problems | -0.83 | 0.45 | -1.85 | 0.149 |
| CBCL Internalizing Problems | -0.96 | 0.27 | -3.52 | **0.002** |
| CBCL Externalizing Problems | -0.23 | 0.29 | -0.80 | 0.607 |
| **State 5** |  |  |  |  |
| Intercept | 39.11 | 10.42 | 3.76 | 0.001 |
| Age | 0.04 | 0.08 | 0.52 | 0.757 |
| Sex | -0.62 | 1.20 | -0.52 | 0.757 |
| Cognition | -0.04 | 0.04 | -1.22 | 0.397 |
| Motion | -19.68 | 1.89 | -10.39 | 2.13E-15 |
| CBCL Attention Problems | -0.89 | 0.24 | -3.72 | **0.001** |
| CBCL Internalizing Problems | -0.67 | 0.14 | -4.64 | **2.44E-05** |
| CBCL Externalizing Problems | -0.13 | 0.15 | -0.86 | 0.589 |
| **State 6** |  |  |  |  |
| Intercept | 1.69 | 9.68 | 0.17 | 0.942 |
| Age | 0.06 | 0.07 | 0.76 | 0.629 |
| Sex | -1.09 | 1.12 | -0.98 | 0.524 |
| Cognition | 0.05 | 0.03 | 1.59 | 0.229 |
| Motion | 16.66 | 1.76 | 9.47 | 2.13E-15 |
| CBCL Attention Problems | -0.72 | 0.22 | -3.28 | **0.004** |
| CBCL Internalizing Problems | -0.22 | 0.13 | -1.62 | 0.219 |
| CBCL Externalizing Problems | -0.14 | 0.14 | -0.99 | 0.524 |
| **State 7** |  |  |  |  |
| Intercept | 10.12 | 15.51 | 0.65 | 0.715 |
| Age | -0.12 | 0.12 | -1.02 | 0.515 |
| Sex | -1.73 | 1.79 | -0.97 | 0.524 |
| Cognition | 0.00 | 0.05 | 0.00 | 0.998 |
| Motion | 71.26 | 2.82 | 25.27 | 2.13E-15 |
| CBCL Attention Problems | 0.22 | 0.35 | 0.61 | 0.730 |
| CBCL Internalizing Problems | 1.04 | 0.21 | 4.88 | **8.91E-06** |
| CBCL Externalizing Problems | -0.21 | 0.22 | -0.94 | 0.540 |
| **State 8** |  |  |  |  |
| Intercept | 5.92 | 9.56 | 0.62 | 0.730 |
| Age | 0.04 | 0.07 | 0.54 | 0.757 |
| Sex | 2.36 | 1.10 | 2.14 | 0.078 |
| Cognition | 0.05 | 0.03 | 1.41 | 0.300 |
| Motion | -8.44 | 1.74 | -4.86 | 9.02E-06 |
| CBCL Attention Problems | 1.21 | 0.22 | 5.54 | **3.44E-07** |
| CBCL Internalizing Problems | 0.14 | 0.13 | 1.06 | 0.497 |
| CBCL Externalizing Problems | 0.36 | 0.14 | 2.62 | **0.026** |
| **State 9** |  |  |  |  |
| Intercept | -9.09 | 6.26 | -1.45 | 0.288 |
| Age | -0.01 | 0.05 | -0.16 | 0.942 |
| Sex | -0.24 | 0.72 | -0.33 | 0.836 |
| Cognition | 0.03 | 0.02 | 1.47 | 0.282 |
| Motion | 29.67 | 1.14 | 26.05 | 2.13E-15 |
| CBCL Attention Problems | 0.77 | 0.14 | 5.39 | **7.22E-07** |
| CBCL Internalizing Problems | 0.44 | 0.09 | 5.13 | **2.67E-06** |
| CBCL Externalizing Problems | 0.08 | 0.09 | 0.86 | 0.589 |
| **State 10** |  |  |  |  |
| Intercept | -11.27 | 22.42 | -0.50 | 0.757 |
| Age | 0.09 | 0.17 | 0.55 | 0.757 |
| Sex | -3.31 | 2.59 | -1.28 | 0.371 |
| Cognition | 0.06 | 0.08 | 0.82 | 0.601 |
| Motion | 60.38 | 4.08 | 14.81 | 2.13E-15 |
| CBCL Attention Problems | 1.55 | 0.51 | 3.02 | **0.008** |
| CBCL Internalizing Problems | 0.96 | 0.31 | 3.09 | **0.007** |
| CBCL Externalizing Problems | 0.11 | 0.32 | 0.35 | 0.836 |
| **State 11** |  |  |  |  |
| Intercept | 34.93 | 19.11 | 1.83 | 0.152 |
| Age | -0.02 | 0.14 | -0.12 | 0.947 |
| Sex | 4.91 | 2.20 | 2.23 | 0.068 |
| Cognition | 0.01 | 0.07 | 0.12 | 0.947 |
| Motion | -35.39 | 3.47 | -10.18 | 2.13E-15 |
| CBCL Attention Problems | 1.87 | 0.44 | 4.28 | **1.04E-04** |
| CBCL Internalizing Problems | 0.93 | 0.26 | 3.51 | **1.71E-03** |
| CBCL Externalizing Problems | 0.65 | 0.28 | 2.35 | 0.052 |
| **State 12** |  |  |  |  |
| Intercept | 44.20 | 16.56 | 2.67 | 0.023 |
| Age | -0.18 | 0.12 | -1.44 | 0.289 |
| Sex | 1.92 | 1.91 | 1.01 | 0.520 |
| Cognition | -0.05 | 0.06 | -0.82 | 0.601 |
| Motion | -13.09 | 3.01 | -4.35 | 8.21E-05 |
| CBCL Attention Problems | 0.44 | 0.38 | 1.17 | 0.422 |
| CBCL Internalizing Problems | 0.41 | 0.23 | 1.79 | 0.161 |
| CBCL Externalizing Problems | 0.98 | 0.24 | 4.09 | **2.24E-04** |

Note: CBCL, Child Behavior Checklist. General Cognition is measured by the NIH Toolbox age corrected scores (28).

**Table S2.** Linear regression models depicting associations between minimum sojourn time and disruptive behavior severity across all latent states accounting for motion

| ***Predictors*** | ***β*** | ***Std. Error*** | ***t*** | ***p_FDR_*** |
| --- | --- | --- | --- | --- |
| **State 1** |  |  |  |  |
| Intercept | -0.338 | 2.322 | -0.145 | 0.96 |
| Age | 0.024 | 0.017 | 1.368 | 0.48 |
| Sex | -0.660 | 0.268 | -2.464 | 0.07 |
| Cognition | -6.40E-05 | 0.008 | -0.008 | 0.99 |
| Motion | 3.638 | 0.422 | 8.616 | 1.56E-15 |
| CBCL Attention Problems | -0.101 | 0.053 | -1.898 | 0.22 |
| CBCL Internalizing Problems | 0.039 | 0.032 | 1.220 | 0.55 |
| CBCL Externalizing Problems | -0.025 | 0.034 | -0.744 | 0.82 |
| **State 2** |  |  |  |  |
| Intercept | 1.909 | 0.964 | 1.981 | 0.21 |
| Age | 0.001 | 0.007 | 0.121 | 0.96 |
| Sex | 0.098 | 0.111 | 0.884 | 0.75 |
| Cognition | 0.001 | 0.003 | 0.439 | 0.93 |
| Motion | -0.444 | 0.175 | -2.536 | 0.07 |
| CBCL Attention Problems | 0.019 | 0.022 | 0.871 | 0.75 |
| CBCL Internalizing Problems | -0.025 | 0.013 | -1.905 | 0.22 |
| CBCL Externalizing Problems | 0.014 | 0.014 | 1.033 | 0.70 |
| **State 3** |  |  |  |  |
| Intercept | 1.914 | 0.481 | 3.982 | 7.12E-04 |
| Age | -0.003 | 0.004 | -0.695 | 0.84 |
| Sex | 0.036 | 0.055 | 0.654 | 0.85 |
| Cognition | -0.001 | 0.002 | -0.430 | 0.93 |
| Motion | -0.489 | 0.087 | -5.591 | 5.80E-07 |
| CBCL Attention Problems | 0.008 | 0.011 | 0.735 | 0.82 |
| CBCL Internalizing Problems | -0.003 | 0.007 | -0.384 | 0.93 |
| CBCL Externalizing Problems | 0.002 | 0.007 | 0.308 | 0.93 |
| **State 4** |  |  |  |  |
| Intercept | 2.014 | 0.452 | 4.451 | 1.33E-04 |
| Age | -0.005 | 0.003 | -1.607 | 0.36 |
| Sex | -0.053 | 0.052 | -1.022 | 0.70 |
| Cognition | 0.001 | 0.002 | 0.360 | 0.93 |
| Motion | -0.101 | 0.082 | -1.227 | 0.55 |
| CBCL Attention Problems | 0.003 | 0.010 | 0.287 | 0.93 |
| CBCL Internalizing Problems | 0.010 | 0.006 | 1.572 | 0.37 |
| CBCL Externalizing Problems | 0.001 | 0.007 | 0.128 | 0.96 |
| **State 5** |  |  |  |  |
| Intercept | 1.984 | 0.711 | 2.791 | 0.04 |
| Age | 0.001 | 0.005 | 0.276 | 0.93 |
| Sex | -0.035 | 0.082 | -0.428 | 0.93 |
| Cognition | -0.001 | 0.002 | -0.335 | 0.93 |
| Motion | -0.504 | 0.129 | -3.898 | 8.35E-04 |
| CBCL Attention Problems | 0.006 | 0.016 | 0.389 | 0.93 |
| CBCL Internalizing Problems | -0.006 | 0.010 | -0.605 | 0.87 |
| CBCL Externalizing Problems | 0.005 | 0.010 | 0.518 | 0.92 |
| **State 6** |  |  |  |  |
| Intercept | 1.506 | 0.311 | 4.844 | 2.41E-05 |
| Age | -0.004 | 0.002 | -1.732 | 0.30 |
| Sex | -0.003 | 0.036 | -0.091 | 0.98 |
| Cognition | -0.001 | 0.001 | -0.644 | 0.85 |
| Motion | 0.332 | 0.057 | 5.867 | 1.51E-07 |
| CBCL Attention Problems | -0.005 | 0.007 | -0.764 | 0.82 |
| CBCL Internalizing Problems | -0.007 | 0.004 | -1.607 | 0.36 |
| CBCL Externalizing Problems | 0.001 | 0.005 | 0.179 | 0.96 |
| **State 7** |  |  |  |  |
| Intercept | 1.839 | 0.439 | 4.188 | 3.56E-04 |
| Age | -0.006 | 0.003 | -1.783 | 0.28 |
| Sex | -0.047 | 0.051 | -0.937 | 0.75 |
| Cognition | 0.002 | 0.001 | 1.038 | 0.70 |
| Motion | -0.160 | 0.080 | -2.002 | 0.21 |
| CBCL Attention Problems | 3.547E-04 | 0.010 | 0.035 | 0.99 |
| CBCL Internalizing Problems | -0.001 | 0.006 | -0.155 | 0.96 |
| CBCL Externalizing Problems | 0.003 | 0.006 | 0.444 | 0.93 |
| **State 8** |  |  |  |  |
| Intercept | 1.918 | 0.723 | 2.655 | 0.05 |
| Age | 0.004 | 0.005 | 0.694 | 0.84 |
| Sex | -0.168 | 0.083 | -2.019 | 0.21 |
| Cognition | 0.002 | 0.002 | 0.856 | 0.75 |
| Motion | -0.548 | 0.131 | -4.171 | 3.56E-04 |
| CBCL Attention Problems | -0.005 | 0.016 | -0.313 | 0.93 |
| CBCL Internalizing Problems | 0.005 | 0.010 | 0.454 | 0.93 |
| CBCL Externalizing Problems | -3.114E-04 | 0.010 | -0.030 | 0.99 |
| **State 9** |  |  |  |  |
| Intercept | 0.216 | 0.460 | 0.469 | 0.93 |
| Age | -0.001 | 0.003 | -0.241 | 0.94 |
| Sex | -0.002 | 0.053 | -0.039 | 0.99 |
| Cognition | 0.002 | 0.002 | 1.329 | 0.51 |
| Motion | 0.695 | 0.084 | 8.309 | 1.18E-14 |
| CBCL Attention Problems | 0.003 | 0.010 | 0.309 | 0.93 |
| CBCL Internalizing Problems | 0.013 | 0.006 | 2.116 | 0.17 |
| CBCL Externalizing Problems | -0.009 | 0.007 | -1.299 | 0.52 |
| **State 10** |  |  |  |  |
| Intercept | 0.589 | 0.585 | 1.007 | 0.70 |
| Age | 0.001 | 0.004 | 0.303 | 0.93 |
| Sex | 0.037 | 0.067 | 0.542 | 0.91 |
| Cognition | 0.001 | 0.002 | 0.579 | 0.89 |
| Motion | 0.096 | 0.106 | 0.903 | 0.75 |
| CBCL Attention Problems | 0.020 | 0.013 | 1.524 | 0.40 |
| CBCL Internalizing Problems | -0.002 | 0.008 | -0.211 | 0.95 |
| CBCL Externalizing Problems | -0.005 | 0.008 | -0.644 | 0.85 |
| **State 11** |  |  |  |  |
| Intercept | 1.092 | 0.285 | 3.832 | 1.01E-03 |
| Age | 0.001 | 0.002 | 0.350 | 0.93 |
| Sex | 0.029 | 0.033 | 0.887 | 0.75 |
| Cognition | -2.712E-04 | 0.001 | -0.279 | 0.93 |
| Motion | -0.203 | 0.052 | -3.917 | 8.35E-04 |
| CBCL Attention Problems | -0.006 | 0.007 | -0.880 | 0.75 |
| CBCL Internalizing Problems | 0.006 | 0.004 | 1.476 | 0.42 |
| CBCL Externalizing Problems | 0.001 | 0.004 | 0.233 | 0.94 |
| **State 12** | -0.338 | 2.322 | -0.145 | 0.96 |
| Intercept | 0.024 | 0.017 | 1.368 | 0.48 |
| Age | -0.660 | 0.268 | -2.464 | 0.07 |
| Sex | -6.396E-05 | 0.008 | -0.008 | 0.99 |
| Cognition | 3.638 | 0.422 | 8.616 | 1.56E-15 |
| Motion | -0.101 | 0.053 | -1.898 | 0.22 |
| CBCL Attention Problems | 0.039 | 0.032 | 1.220 | 0.55 |
| CBCL Internalizing Problems | -0.025 | 0.034 | -0.744 | 0.82 |
| CBCL Externalizing Problems | -0.338 | 2.322 | -0.145 | 0.96 |

Note: CBCL, Child Behavior Checklist. General Cognition is measured by the NIH Toolbox age corrected scores (28).

**Table S3.** Linear regression models depicting associations between maximum sojourn time and disruptive behavior severity across all latent states accounting for motion

| ***Predictors*** | ***β*** | ***Std. Error*** | ***t*** | ***p_FDR_*** |
| --- | --- | --- | --- | --- |
| **State 1** |  |  |  |  |
| Intercept | 5.58 | 5.74 | 0.97 | 0.64 |
| Age | -0.02 | 0.04 | -0.47 | 0.81 |
| Sex | -0.98 | 0.66 | -1.48 | 0.36 |
| Cognition | -9.72E-04 | 0.02 | -0.05 | 0.98 |
| Motion | 10.49 | 1.04 | 10.06 | 2.23E-21 |
| CBCL Attention Problems | -0.25 | 0.13 | -1.91 | 0.21 |
| CBCL Internalizing Problems | 0.03 | 0.08 | 0.33 | 0.86 |
| CBCL Externalizing Problems | 0.04 | 0.08 | 0.49 | 0.81 |
| **State 2** |  |  |  |  |
| Intercept | 19.81 | 5.87 | 3.37 | 4.37E-03 |
| Age | -0.02 | 0.04 | -0.37 | 0.85 |
| Sex | 1.15 | 0.68 | 1.69 | 0.28 |
| Cognition | 0.01 | 0.02 | 0.55 | 0.81 |
| Motion | -10.90 | 1.07 | -10.21 | 0.00 |
| CBCL Attention Problems | -0.18 | 0.13 | -1.33 | 0.42 |
| CBCL Internalizing Problems | -0.12 | 0.08 | -1.43 | 0.37 |
| CBCL Externalizing Problems | -0.06 | 0.09 | -0.70 | 0.77 |
| **State 3** |  |  |  |  |
| Intercept | 20.50 | 5.85 | 3.51 | 2.87E-03 |
| Age | -0.03 | 0.04 | -0.69 | 0.77 |
| Sex | 0.58 | 0.67 | 0.86 | 0.67 |
| Cognition | -0.03 | 0.02 | -1.70 | 0.28 |
| Motion | -8.67 | 1.06 | -8.15 | 1.49E-14 |
| CBCL Attention Problems | -0.10 | 0.13 | -0.72 | 0.77 |
| CBCL Internalizing Problems | -0.10 | 0.08 | -1.30 | 0.42 |
| CBCL Externalizing Problems | -0.08 | 0.08 | -0.92 | 0.64 |
| **State 4** |  |  |  |  |
| Intercept | 10.84 | 3.82 | 2.84 | 0.02 |
| Age | 0.01 | 0.03 | 0.31 | 0.86 |
| Sex | 0.04 | 0.44 | 0.08 | 0.98 |
| Cognition | 3.82E-03 | 0.01 | 0.29 | 0.86 |
| Motion | -4.27 | 0.69 | -6.15 | 1.12E-08 |
| CBCL Attention Problems | -0.04 | 0.09 | -0.40 | 0.84 |
| CBCL Internalizing Problems | -0.14 | 0.05 | -2.61 | **0.04** |
| CBCL Externalizing Problems | 0.06 | 0.06 | 1.00 | 0.64 |
| **State 5** |  |  |  |  |
| Intercept | 9.08 | 2.06 | 4.41 | 7.84E-05 |
| Age | -0.01 | 0.02 | -0.61 | 0.81 |
| Sex | 0.29 | 0.24 | 1.20 | 0.49 |
| Cognition | 2.85E-05 | 0.01 | 0.00 | 1.00 |
| Motion | -3.81 | 0.37 | -10.17 | 1.14E-21 |
| CBCL Attention Problems | -3.82E-03 | 0.05 | -0.08 | 0.98 |
| CBCL Internalizing Problems | -0.09 | 0.03 | -3.17 | **0.01** |
| CBCL Externalizing Problems | -0.02 | 0.03 | -0.58 | 0.81 |
| **State 6** |  |  |  |  |
| Intercept | -1.14 | 2.34 | -0.48 | 0.81 |
| Age | 0.02 | 0.02 | 1.41 | 0.37 |
| Sex | -0.26 | 0.27 | -0.95 | 0.64 |
| Cognition | 0.01 | 0.01 | 1.43 | 0.37 |
| Motion | 3.85 | 0.43 | 9.03 | 1.48E-17 |
| CBCL Attention Problems | -0.08 | 0.05 | -1.41 | 0.37 |
| CBCL Internalizing Problems | 0.01 | 0.03 | 0.29 | 0.86 |
| CBCL Externalizing Problems | -3.13E-03 | 0.03 | -0.09 | 0.98 |
| **State 7** |  |  |  |  |
| Intercept | 3.45 | 5.71 | 0.60 | 0.81 |
| Age | -0.01 | 0.04 | -0.34 | 0.86 |
| Sex | -0.69 | 0.66 | -1.05 | 0.61 |
| Cognition | -0.02 | 0.02 | -0.93 | 0.64 |
| Motion | 14.40 | 1.04 | 13.88 | 0.00 |
| CBCL Attention Problems | 1.64E-03 | 0.13 | 0.01 | 1.00 |
| CBCL Internalizing Problems | 0.39 | 0.08 | 4.95 | **6.53E-06** |
| CBCL Externalizing Problems | -0.02 | 0.08 | -0.24 | 0.90 |
| **State 8** |  |  |  |  |
| Intercept | 6.30 | 2.30 | 2.74 | 0.03 |
| Age | -0.01 | 0.02 | -0.46 | 0.81 |
| Sex | 0.12 | 0.27 | 0.46 | 0.81 |
| Cognition | 0.01 | 0.01 | 1.72 | 0.28 |
| Motion | -2.25 | 0.42 | -5.36 | 8.38E-07 |
| CBCL Attention Problems | 0.17 | 0.05 | 3.15 | **0.01** |
| CBCL Internalizing Problems | 0.02 | 0.03 | 0.76 | 0.74 |
| CBCL Externalizing Problems | 0.03 | 0.03 | 0.94 | 0.64 |
| **State 9** |  |  |  |  |
| Intercept | -0.74 | 1.69 | -0.44 | 0.81 |
| Age | -0.01 | 0.01 | -0.51 | 0.81 |
| Sex | 0.10 | 0.19 | 0.51 | 0.81 |
| Cognition | 0.01 | 0.01 | 1.04 | 0.61 |
| Motion | 7.00 | 0.31 | 22.83 | 9.58E-89 |
| CBCL Attention Problems | 0.07 | 0.04 | 1.74 | 0.28 |
| CBCL Internalizing Problems | 0.09 | 0.02 | 3.68 | **1.57E-03** |
| CBCL Externalizing Problems | -0.01 | 0.02 | -0.49 | 0.81 |
| **State 10** |  |  |  |  |
| Intercept | -4.51 | 5.08 | -0.89 | 0.65 |
| Age | 0.05 | 0.04 | 1.30 | 0.42 |
| Sex | -0.38 | 0.59 | -0.65 | 0.80 |
| Cognition | 0.03 | 0.02 | 1.63 | 0.31 |
| Motion | 5.43 | 0.92 | 5.88 | 0.00 |
| CBCL Attention Problems | 0.20 | 0.12 | 1.72 | 0.28 |
| CBCL Internalizing Problems | 0.11 | 0.07 | 1.59 | 0.32 |
| CBCL Externalizing Problems | -0.04 | 0.07 | -0.59 | 0.81 |
| **State 11** |  |  |  |  |
| Intercept | 13.80 | 4.96 | 2.78 | 0.03 |
| Age | -0.03 | 0.04 | -0.80 | 0.71 |
| Sex | 0.88 | 0.57 | 1.54 | 0.34 |
| Cognition | 3.68E-03 | 0.02 | 0.22 | 0.90 |
| Motion | -6.85 | 0.90 | -7.59 | 8.75E-13 |
| CBCL Attention Problems | 0.27 | 0.11 | 2.34 | 0.08 |
| CBCL Internalizing Problems | -0.01 | 0.07 | -0.14 | 0.96 |
| CBCL Externalizing Problems | 0.11 | 0.07 | 1.53 | 0.34 |
| **State 12** |  |  |  |  |
| Intercept | 5.58 | 5.74 | 0.97 | 0.64 |
| Age | -0.02 | 0.04 | -0.47 | 0.81 |
| Sex | -0.98 | 0.66 | -1.48 | 0.36 |
| Cognition | -9.72E-04 | 0.02 | -0.05 | 0.98 |
| Motion | 10.49 | 1.04 | 10.06 | 2.23E-21 |
| CBCL Attention Problems | -0.25 | 0.13 | -1.91 | 0.21 |
| CBCL Internalizing Problems | 0.03 | 0.08 | 0.33 | 0.86 |
| CBCL Externalizing Problems | 0.04 | 0.08 | 0.49 | 0.81 |

Note: CBCL, Child Behavior Checklist. General Cognition is measured by the NIH Toolbox age corrected scores (28).

**Table S4.** Graph Theory Metrics Quantifying Brain-Wide Topology for Each of the 12 States

| **Brain State** | **Betweenness** | **Centrality** | **Local Efficiency** | **Global Efficiency** | **Clustering Coefficient** |
| --- | --- | --- | --- | --- | --- |
| State 1 | 18.970 | 8.424 | 0.352 | 0.223 | 0.260 |
| State 2 | 40.848 | 6.182 | 0.321 | 0.213 | 0.250 |
| State 3 | 37.394 | 5.576 | 0.292 | 0.191 | 0.216 |
| State 4 | 17.576 | 10.848 | 0.365 | 0.263 | 0.281 |
| State 5 | 37.879 | 5.758 | 0.338 | 0.213 | 0.254 |
| State 6 | 16.364 | 12.364 | 0.380 | 0.213 | 0.298 |
| State 7 | 18.242 | 9.152 | 0.354 | 0.230 | 0.258 |
| State 8 | 40.303 | 5.394 | 0.338 | 0.205 | 0.239 |
| State 9 | 16.121 | 12.667 | 0.388 | 0.281 | 0.310 |
| State 10 | 16.545 | 11.697 | 0.368 | 0.269 | 0.291 |
| State 11 | 34.364 | 5.697 | 0.314 | 0.191 | 0.238 |
| State 12 | 40.121 | 5.212 | 0.297 | 0.185 | 0.221 |

**Table S5.** Exploratory Follow-up Analysis of Graph Theory Metrics Related to Intrinsic Networks and Brain State

| **Network** | **State** | **Betweenness** | | | | **Degree** | | | **Local Efficiency** | | | **Clustering Coefficient** | |
| --- | --- | --- | --- | --- | --- | --- | --- | --- | --- | --- | --- | --- | --- |
|  |  | **Mean (SD)** | **Median** |  | **Mean (SD)** | | **Median** |  | **Mean (SD)** | **Median** |  | **Mean (SD)** | **Median** |
| **VIS** | State 1 | 10 (19.08) | 2.00 |  | 6.8 (2.49) | | 7.00 |  | 0.48 (0.03) | 0.47 |  | 0.42 (0.08) | 0.40 |
|  | State 2 | 12 (13.27) | 6.00 |  | 5.4 (1.67) | | 5.00 |  | 0.5 (0.03) | 0.49 |  | 0.4 (0.09) | 0.38 |
|  | State 3 | 12.8 (18.9) | 8.00 |  | 5.2 (1.48) | | 5.00 |  | 0.45 (0.02) | 0.46 |  | 0.34 (0.08) | 0.31 |
|  | State 4 | 2 (3.46) | 0.00 |  | 8.2 (2.59) | | 9.00 |  | 0.48 (0.01) | 0.48 |  | 0.43 (0.03) | 0.42 |
|  | State 5 | 13.6 (25.04) | 2.00 |  | 5.4 (1.82) | | 5.00 |  | 0.51 (0.04) | 0.51 |  | 0.42 (0.1) | 0.43 |
|  | State 6 | 12 (14.7) | 4.00 |  | 9.4 (2.61) | | 10.00 |  | 0.46 (0.02) | 0.47 |  | 0.38 (0.05) | 0.40 |
|  | State 7 | 11.2 (15.79) | 6.00 |  | 6.8 (2.59) | | 7.00 |  | 0.48 (0.04) | 0.47 |  | 0.39 (0.09) | 0.37 |
|  | State 8 | 16.8 (27.15) | 2.00 |  | 5.4 (1.82) | | 5.00 |  | 0.49 (0.04) | 0.50 |  | 0.38 (0.1) | 0.38 |
|  | State 9 | 11.6 (15.58) | 2.00 |  | 9.6 (2.7) | | 10.00 |  | 0.48 (0.02) | 0.48 |  | 0.4 (0.04) | 0.42 |
|  | State 10 | 2.4 (1.67) | 2.00 |  | 8.2 (2.59) | | 9.00 |  | 0.49 (0.01) | 0.49 |  | 0.43 (0.04) | 0.41 |
|  | State 11 | 11.2 (19.58) | 2.00 |  | 5.4 (1.67) | | 5.00 |  | 0.49 (0.02) | 0.50 |  | 0.39 (0.08) | 0.38 |
|  | State 12 | 24.8 (32.94) | 12.00 |  | 4.8 (1.3) | | 5.00 |  | 0.44 (0.08) | 0.46 |  | 0.34 (0.12) | 0.28 |
| **SM** | State 1 | 4.5 (6.61) | 2.00 |  | 7 (1.83) | | 7.00 |  | 0.44 (0.01) | 0.45 |  | 0.38 (0.06) | 0.38 |
|  | State 2 | 12.5 (14.46) | 12.00 |  | 7.75 (1.26) | | 8.00 |  | 0.42 (0.01) | 0.41 |  | 0.36 (0.04) | 0.35 |
|  | State 3 | 9.5 (12.37) | 6.00 |  | 6.75 (1.5) | | 7.00 |  | 0.41 (0.01) | 0.41 |  | 0.34 (0.04) | 0.32 |
|  | State 4 | 12 (9.93) | 11.00 |  | 11 (3.56) | | 12.00 |  | 0.42 (0.01) | 0.42 |  | 0.35 (0.04) | 0.35 |
|  | State 5 | 0 (0) | 0.00 |  | 2.5 (0.58) | | 2.50 |  | 0.42 (0.04) | 0.42 |  | 0.42 (0.04) | 0.42 |
|  | State 6 | 12 (7.66) | 10.00 |  | 12.75 (1.89) | | 13.50 |  | 0.43 (0.01) | 0.43 |  | 0.36 (0.02) | 0.36 |
|  | State 7 | 4.5 (9) | 0.00 |  | 7.5 (2.38) | | 7.50 |  | 0.44 (0.01) | 0.45 |  | 0.38 (0.04) | 0.39 |
|  | State 8 | 2.5 (5) | 0.00 |  | 2.5 (1.29) | | 2.50 |  | 0.32 (0.22) | 0.39 |  | 0.29 (0.22) | 0.34 |
|  | State 9 | 18 (17.96) | 11.00 |  | 13.5 (1.73) | | 13.00 |  | 0.43 (0.02) | 0.44 |  | 0.37 (0.04) | 0.38 |
|  | State 10 | 8 (4.9) | 8.00 |  | 11.75 (2.87) | | 12.50 |  | 0.43 (0) | 0.43 |  | 0.37 (0.03) | 0.36 |
|  | State 11 | 12.5 (15) | 10.00 |  | 7.5 (1.29) | | 7.50 |  | 0.41 (0.01) | 0.41 |  | 0.33 (0.04) | 0.33 |
|  | State 12 | 10.5 (19.69) | 1.00 |  | 4.75 (1.71) | | 4.50 |  | 0.4 (0.02) | 0.40 |  | 0.35 (0.08) | 0.36 |
| **DAN** | State 1 | 38.4 (52.22) | 14.00 |  | 13.2 (3.56) | | 12.00 |  | 0.41 (0.03) | 0.41 |  | 0.25 (0.05) | 0.26 |
|  | State 2 | 29.6 (19.15) | 22.00 |  | 9.2 (1.3) | | 9.00 |  | 0.4 (0.02) | 0.41 |  | 0.25 (0.04) | 0.26 |
|  | State 3 | 10 (10.86) | 6.00 |  | 8 (1) | | 8.00 |  | 0.39 (0.02) | 0.38 |  | 0.24 (0.03) | 0.24 |
|  | State 4 | 30 (42.78) | 12.00 |  | 16.2 (4.21) | | 15.00 |  | 0.44 (0.03) | 0.44 |  | 0.3 (0.05) | 0.31 |
|  | State 5 | 32.4 (21.09) | 32.00 |  | 9 (1) | | 9.00 |  | 0.4 (0.01) | 0.40 |  | 0.25 (0.02) | 0.24 |
|  | State 6 | 32.8 (36.65) | 18.00 |  | 19.6 (3.13) | | 19.00 |  | 0.43 (0.02) | 0.44 |  | 0.31 (0.04) | 0.32 |
|  | State 7 | 33.2 (12.21) | 34.00 |  | 15.2 (1.79) | | 14.00 |  | 0.41 (0.01) | 0.41 |  | 0.26 (0.02) | 0.27 |
|  | State 8 | 32.4 (19.62) | 28.00 |  | 8.4 (1.14) | | 8.00 |  | 0.4 (0.02) | 0.40 |  | 0.26 (0.04) | 0.24 |
|  | State 9 | 30.8 (38.38) | 16.00 |  | 20 (2.65) | | 19.00 |  | 0.44 (0.02) | 0.44 |  | 0.32 (0.04) | 0.33 |
|  | State 10 | 30.4 (34.71) | 18.00 |  | 17.8 (3.03) | | 17.00 |  | 0.43 (0.01) | 0.44 |  | 0.31 (0.03) | 0.31 |
|  | State 11 | 19.6 (18.73) | 18.00 |  | 8.4 (0.89) | | 9.00 |  | 0.41 (0.02) | 0.42 |  | 0.25 (0.03) | 0.27 |
|  | State 12 | 34.8 (12.21) | 32.00 |  | 8.4 (1.14) | | 8.00 |  | 0.37 (0.01) | 0.36 |  | 0.21 (0.03) | 0.20 |
| **VAN** | State 1 | 6.5 (3) | 8.00 |  | 7.25 (0.96) | | 7.50 |  | 0.4 (0.02) | 0.40 |  | 0.31 (0.01) | 0.32 |
|  | State 2 | 15.5 (16.84) | 15.00 |  | 6 (2.94) | | 5.50 |  | 0.37 (0.09) | 0.39 |  | 0.32 (0.12) | 0.33 |
|  | State 3 | 13 (16.53) | 8.00 |  | 5.25 (2.06) | | 5.00 |  | 0.35 (0.08) | 0.37 |  | 0.29 (0.11) | 0.28 |
|  | State 4 | 7.5 (9.98) | 4.00 |  | 9.75 (2.06) | | 10.00 |  | 0.43 (0.03) | 0.43 |  | 0.34 (0.02) | 0.34 |
|  | State 5 | 21.5 (24.19) | 19.00 |  | 5.75 (1.89) | | 6.50 |  | 0.38 (0.07) | 0.39 |  | 0.32 (0.12) | 0.32 |
|  | State 6 | 3.5 (3.42) | 3.00 |  | 9.75 (1.71) | | 9.50 |  | 0.44 (0.02) | 0.44 |  | 0.37 (0.01) | 0.37 |
|  | State 7 | 8.5 (6.61) | 9.00 |  | 8.5 (1.73) | | 8.50 |  | 0.4 (0.02) | 0.41 |  | 0.3 (0.04) | 0.29 |
|  | State 8 | 11 (17.09) | 4.00 |  | 5.25 (2.36) | | 6.00 |  | 0.41 (0.06) | 0.39 |  | 0.34 (0.14) | 0.34 |
|  | State 9 | 1.5 (3) | 0.00 |  | 9.25 (2.06) | | 9.00 |  | 0.45 (0.02) | 0.46 |  | 0.39 (0.02) | 0.39 |
|  | State 10 | 6.5 (4.73) | 8.00 |  | 10.5 (2.65) | | 11.00 |  | 0.44 (0.03) | 0.43 |  | 0.36 (0.03) | 0.35 |
|  | State 11 | 11.5 (13.3) | 8.00 |  | 5.5 (2.08) | | 5.50 |  | 0.37 (0.09) | 0.38 |  | 0.31 (0.11) | 0.30 |
|  | State 12 | 12 (15.32) | 8.00 |  | 5.5 (2.08) | | 5.50 |  | 0.36 (0.08) | 0.36 |  | 0.3 (0.12) | 0.27 |
| **LIMB** | State 1 | 0 (0) | 0.00 |  | 0.5 (0.71) | | 0.50 |  | 0 (0) | 0.00 |  | 0 (0) | 0.00 |
|  | State 2 | 0 (0) | 0.00 |  | 1.5 (0.71) | | 1.50 |  | 0.23 (0.33) | 0.23 |  | 0.23 (0.33) | 0.23 |
|  | State 3 | 0 (0) | 0.00 |  | 1.5 (0.71) | | 1.50 |  | 0.22 (0.31) | 0.22 |  | 0.22 (0.31) | 0.22 |
|  | State 4 | 0 (0) | 0.00 |  | 0.5 (0.71) | | 0.50 |  | 0 (0) | 0.00 |  | 0 (0) | 0.00 |
|  | State 5 | 0 (0) | 0.00 |  | 1.5 (0.71) | | 1.50 |  | 0.23 (0.33) | 0.23 |  | 0.23 (0.33) | 0.23 |
|  | State 6 | 0 (0) | 0.00 |  | 0 (0) | | 0.00 |  | 0 (0) | 0.00 |  | 0 (0) | 0.00 |
|  | State 7 | 0 (0) | 0.00 |  | 0.5 (0.71) | | 0.50 |  | 0 (0) | 0.00 |  | 0 (0) | 0.00 |
|  | State 8 | 0 (0) | 0.00 |  | 1.5 (0.71) | | 1.50 |  | 0.24 (0.34) | 0.24 |  | 0.24 (0.34) | 0.24 |
|  | State 9 | 0 (0) | 0.00 |  | 0 (0) | | 0.00 |  | 0 (0) | 0.00 |  | 0 (0) | 0.00 |
|  | State 10 | 0 (0) | 0.00 |  | 0.5 (0.71) | | 0.50 |  | 0 (0) | 0.00 |  | 0 (0) | 0.00 |
|  | State 11 | 0 (0) | 0.00 |  | 1 (1.41) | | 1.00 |  | 0.24 (0.33) | 0.24 |  | 0.24 (0.33) | 0.24 |
|  | State 12 | 0 (0) | 0.00 |  | 1.5 (0.71) | | 1.50 |  | 0.22 (0.31) | 0.22 |  | 0.22 (0.31) | 0.22 |
| **CON** | State 1 | 18 (9.17) | 16.00 |  | 12.67 (1.53) | | 13.00 |  | 0.4 (0.02) | 0.39 |  | 0.25 (0.01) | 0.25 |
|  | State 2 | 78.67 (74.57) | 68.00 |  | 6.33 (4.16) | | 5.00 |  | 0.23 (0.13) | 0.16 |  | 0.15 (0.03) | 0.14 |
|  | State 3 | 62 (67.44) | 46.00 |  | 5.67 (3.06) | | 5.00 |  | 0.22 (0.12) | 0.16 |  | 0.14 (0.02) | 0.14 |
|  | State 4 | 32.67 (20.82) | 26.00 |  | 15.67 (2.89) | | 14.00 |  | 0.42 (0.01) | 0.42 |  | 0.28 (0.02) | 0.29 |
|  | State 5 | 97.33 (64.38) | 82.00 |  | 8 (4) | | 8.00 |  | 0.29 (0.08) | 0.33 |  | 0.16 (0.01) | 0.16 |
|  | State 6 | 12 (7.21) | 14.00 |  | 17.33 (2.52) | | 17.00 |  | 0.44 (0) | 0.44 |  | 0.33 (0.02) | 0.34 |
|  | State 7 | 26.67 (27.01) | 26.00 |  | 13.67 (4.51) | | 14.00 |  | 0.4 (0.02) | 0.40 |  | 0.25 (0.03) | 0.26 |
|  | State 8 | 87.33 (67.66) | 62.00 |  | 7.67 (3.79) | | 6.00 |  | 0.35 (0.03) | 0.35 |  | 0.17 (0.02) | 0.17 |
|  | State 9 | 18.67 (11.37) | 22.00 |  | 18.67 (2.08) | | 18.00 |  | 0.44 (0) | 0.44 |  | 0.33 (0.02) | 0.34 |
|  | State 10 | 22.67 (18.9) | 16.00 |  | 17 (3) | | 17.00 |  | 0.42 (0) | 0.43 |  | 0.31 (0.02) | 0.32 |
|  | State 11 | 56 (53.03) | 54.00 |  | 6 (3.61) | | 5.00 |  | 0.23 (0.15) | 0.16 |  | 0.16 (0.04) | 0.13 |
|  | State 12 | 52.67 (49.08) | 56.00 |  | 5.33 (2.52) | | 5.00 |  | 0.22 (0.13) | 0.16 |  | 0.15 (0.02) | 0.14 |
| **DMN** | State 1 | 50.8 (61.4) | 22.00 |  | 12.2 (6.76) | | 10.00 |  | 0.36 (0.04) | 0.38 |  | 0.23 (0.06) | 0.25 |
|  | State 2 | 103.6 (86.3) | 82.00 |  | 7 (6.24) | | 4.00 |  | 0.23 (0.14) | 0.17 |  | 0.16 (0.08) | 0.15 |
|  | State 3 | 114.4 (127.9) | 66.00 |  | 7 (6.2) | | 5.00 |  | 0.2 (0.08) | 0.19 |  | 0.12 (0.03) | 0.12 |
|  | State 4 | 31.6 (26.74) | 22.00 |  | 14.8 (5.45) | | 13.00 |  | 0.4 (0.03) | 0.41 |  | 0.28 (0.04) | 0.30 |
|  | State 5 | 97.2 (96.46) | 70.00 |  | 8.2 (5.54) | | 6.00 |  | 0.31 (0.07) | 0.32 |  | 0.15 (0.06) | 0.12 |
|  | State 6 | 27.6 (18.62) | 18.00 |  | 17.4 (4.77) | | 16.00 |  | 0.41 (0.01) | 0.41 |  | 0.29 (0.02) | 0.29 |
|  | State 7 | 42.8 (52.57) | 26.00 |  | 12.6 (6.02) | | 11.00 |  | 0.38 (0.04) | 0.39 |  | 0.26 (0.06) | 0.24 |
|  | State 8 | 122 (139.03) | 84.00 |  | 7.8 (5.85) | | 6.00 |  | 0.34 (0.06) | 0.34 |  | 0.18 (0.08) | 0.16 |
|  | State 9 | 22.4 (19.36) | 24.00 |  | 17.6 (4.83) | | 17.00 |  | 0.42 (0.01) | 0.42 |  | 0.31 (0.02) | 0.32 |
|  | State 10 | 36 (27.02) | 20.00 |  | 16.6 (5.22) | | 16.00 |  | 0.4 (0.03) | 0.41 |  | 0.29 (0.04) | 0.29 |
|  | State 11 | 97.2 (113.39) | 62.00 |  | 6.6 (6.43) | | 4.00 |  | 0.26 (0.12) | 0.21 |  | 0.17 (0.07) | 0.16 |
|  | State 12 | 100.8 (107.8) | 66.00 |  | 6.6 (5.32) | | 5.00 |  | 0.28 (0.11) | 0.30 |  | 0.18 (0.09) | 0.16 |
| **CERBLM** | State 1 | 1.5 (3) | 0.00 |  | 2.5 (3.7) | | 1.00 |  | 0.1 (0.2) | 0.00 |  | 0.07 (0.14) | 0.00 |
|  | State 2 | 31.5 (42.38) | 16.00 |  | 2.75 (2.06) | | 3.00 |  | 0.08 (0.06) | 0.10 |  | 0.08 (0.06) | 0.10 |
|  | State 3 | 19 (38) | 0.00 |  | 1.5 (2.38) | | 0.50 |  | 0.01 (0.02) | 0.00 |  | 0.01 (0.02) | 0.00 |
|  | State 4 | 2 (4) | 0.00 |  | 3.5 (5) | | 1.00 |  | 0.1 (0.21) | 0.00 |  | 0.08 (0.17) | 0.00 |
|  | State 5 | 12 (9.09) | 13.00 |  | 2.75 (2.5) | | 2.50 |  | 0.09 (0.12) | 0.06 |  | 0.06 (0.07) | 0.06 |
|  | State 6 | 1 (2) | 0.00 |  | 4.5 (7.05) | | 1.50 |  | 0.21 (0.24) | 0.21 |  | 0.19 (0.22) | 0.17 |
|  | State 7 | 1.5 (3) | 0.00 |  | 3 (4.69) | | 1.00 |  | 0.1 (0.2) | 0.00 |  | 0.06 (0.12) | 0.00 |
|  | State 8 | 21.5 (43) | 0.00 |  | 2 (2.71) | | 1.00 |  | 0.06 (0.12) | 0.00 |  | 0.03 (0.06) | 0.00 |
|  | State 9 | 3 (6) | 0.00 |  | 5 (8.04) | | 1.50 |  | 0.22 (0.25) | 0.21 |  | 0.19 (0.23) | 0.17 |
|  | State 10 | 0.5 (1) | 0.00 |  | 3.75 (5.5) | | 1.00 |  | 0.11 (0.21) | 0.00 |  | 0.09 (0.18) | 0.00 |
|  | State 11 | 25.5 (51) | 0.00 |  | 1.5 (2.38) | | 0.50 |  | 0.01 (0.02) | 0.00 |  | 0.01 (0.02) | 0.00 |
|  | State 12 | 22.5 (45) | 0.00 |  | 1.5 (2.38) | | 0.50 |  | 0.01 (0.02) | 0.00 |  | 0.01 (0.02) | 0.00 |
| **TP** | State 1 | 26 (NA) | 26.00 |  | 11 (NA) | | 11.00 |  | 0.38 (NA) | 0.38 |  | 0.24 (NA) | 0.24 |
|  | State 2 | 148 (NA) | 148.00 |  | 8 (NA) | | 8.00 |  | 0.29 (NA) | 0.29 |  | 0.23 (NA) | 0.23 |
|  | State 3 | 196 (NA) | 196.00 |  | 9 (NA) | | 9.00 |  | 0.23 (NA) | 0.23 |  | 0.17 (NA) | 0.17 |
|  | State 4 | 78 (NA) | 78.00 |  | 17 (NA) | | 17.00 |  | 0.35 (NA) | 0.35 |  | 0.24 (NA) | 0.24 |
|  | State 5 | 108 (NA) | 108.00 |  | 6 (NA) | | 6.00 |  | 0.18 (NA) | 0.18 |  | 0.14 (NA) | 0.14 |
|  | State 6 | 76 (NA) | 76.00 |  | 16 (NA) | | 16.00 |  | 0.36 (NA) | 0.36 |  | 0.26 (NA) | 0.26 |
|  | State 7 | 28 (NA) | 28.00 |  | 11 (NA) | | 11.00 |  | 0.39 (NA) | 0.39 |  | 0.26 (NA) | 0.26 |
|  | State 8 | 72 (NA) | 72.00 |  | 5 (NA) | | 5.00 |  | 0.34 (NA) | 0.34 |  | 0.16 (NA) | 0.16 |
|  | State 9 | 62 (NA) | 62.00 |  | 15 (NA) | | 15.00 |  | 0.37 (NA) | 0.37 |  | 0.3 (NA) | 0.30 |
|  | State 10 | 74 (NA) | 74.00 |  | 17 (NA) | | 17.00 |  | 0.37 (NA) | 0.37 |  | 0.26 (NA) | 0.26 |
|  | State 11 | 128 (NA) | 128.00 |  | 8 (NA) | | 8.00 |  | 0.28 (NA) | 0.28 |  | 0.21 (NA) | 0.21 |
|  | State 12 | 184 (NA) | 184.00 |  | 7 (NA) | | 7.00 |  | 0.19 (NA) | 0.19 |  | 0.13 (NA) | 0.13 |

Note: CERBLM, cerebellar; CON, Frontoparietal control network; DMN, default mode network; DAN, Dorsal Attention Network; LIMB, limbic; VIS, visual network; SM, somatomotor; TP, temporo parietal; and VAN, Ventral Attention Network.

**Table S6.** Replication of linear regression models depicting associations between occupancy

time and disruptive behavior severity across all latent states for resting-state runs 3-4

| ***Predictors*** | ***β*** | ***Std. Error*** | ***t*** | ***p _FDR_*** |
| --- | --- | --- | --- | --- |
| **State 1** |  |  |  |  |
| Intercept | 52.09 | 37.95 | 1.37 | 0.409 |
| Age | -0.23 | 0.29 | -0.79 | 0.711 |
| Sex | 4.28 | 4.38 | 0.98 | 0.617 |
| Cognition | -0.19 | 0.13 | -1.43 | 0.402 |
| Motion | 117.54 | 6.90 | 17.03 | 4.80E-15 |
| CBCL Attention Problems | 2.29 | 0.87 | 2.65 | **0.047** |
| CBCL Internalizing Problems | 0.98 | 0.52 | 1.88 | 0.208 |
| CBCL Externalizing Problems | 0.75 | 0.55 | 1.37 | 0.409 |
| **State 2** |  |  |  |  |
| Intercept | -0.10 | 5.19 | -0.02 | 0.991 |
| Age | 0.01 | 0.04 | 0.29 | 0.904 |
| Sex | -0.03 | 0.60 | -0.05 | 0.991 |
| Cognition | 0.00 | 0.02 | 0.02 | 0.991 |
| Motion | -1.88 | 0.94 | -2.00 | 0.170 |
| CBCL Attention Problems | 0.36 | 0.12 | 3.05 | **0.017** |
| CBCL Internalizing Problems | 0.01 | 0.07 | 0.18 | 0.959 |
| CBCL Externalizing Problems | 0.03 | 0.08 | 0.37 | 0.880 |
| **State 3** |  |  |  |  |
| Intercept | -7.90 | 4.10 | -1.93 | 0.192 |
| Age | 0.07 | 0.03 | 2.24 | 0.116 |
| Sex | 0.23 | 0.47 | 0.48 | 0.835 |
| Cognition | 0.00 | 0.01 | -0.18 | 0.959 |
| Motion | 0.01 | 0.74 | 0.01 | 0.991 |
| CBCL Attention Problems | 0.20 | 0.09 | 2.13 | 0.135 |
| CBCL Internalizing Problems | -0.02 | 0.06 | -0.37 | 0.880 |
| CBCL Externalizing Problems | -0.02 | 0.06 | -0.36 | 0.880 |
| **State 4** |  |  |  |  |
| Intercept | 10.63 | 9.44 | 1.13 | 0.549 |
| Age | -0.04 | 0.07 | -0.55 | 0.822 |
| Sex | -0.71 | 1.09 | -0.65 | 0.788 |
| Cognition | -0.02 | 0.03 | -0.67 | 0.788 |
| Motion | -0.13 | 1.72 | -0.07 | 0.991 |
| CBCL Attention Problems | 0.36 | 0.22 | 1.68 | 0.301 |
| CBCL Internalizing Problems | -0.04 | 0.13 | -0.34 | 0.880 |
| CBCL Externalizing Problems | 0.21 | 0.14 | 1.55 | 0.342 |
| **State 5** |  |  |  |  |
| Intercept | 6.77 | 14.46 | 0.47 | 0.835 |
| Age | -0.13 | 0.11 | -1.22 | 0.507 |
| Sex | -2.30 | 1.67 | -1.38 | 0.409 |
| Cognition | 0.11 | 0.05 | 2.16 | 0.130 |
| Motion | 43.55 | 2.63 | 16.57 | 4.80E-15 |
| CBCL Attention Problems | 0.77 | 0.33 | 2.33 | 0.10 |
| CBCL Internalizing Problems | -0.01 | 0.20 | -0.06 | 0.991 |
| CBCL Externalizing Problems | 0.60 | 0.21 | 2.86 | **0.029** |
| **State 6** |  |  |  |  |
| Intercept | -6.90 | 19.86 | -0.35 | 0.880 |
| Age | 0.15 | 0.15 | 0.99 | 0.617 |
| Sex | 1.85 | 2.29 | 0.81 | 0.707 |
| Cognition | -0.07 | 0.07 | -1.06 | 0.576 |
| Motion | 64.67 | 3.61 | 17.91 | 4.80E-15 |
| CBCL Attention Problems | 1.16 | 0.45 | 2.55 | 0.055 |
| CBCL Internalizing Problems | -0.01 | 0.27 | -0.03 | 0.991 |
| CBCL Externalizing Problems | 0.78 | 0.29 | 2.73 | **0.039** |
| **State 7** |  |  |  |  |
| Intercept | 181.41 | 54.40 | 3.34 | 0.008 |
| Age | 0.44 | 0.41 | 1.07 | 0.576 |
| Sex | -0.29 | 6.27 | -0.05 | 0.991 |
| Cognition | 0.11 | 0.19 | 0.58 | 0.822 |
| Motion | -164.68 | 9.89 | -16.65 | 4.80E-15 |
| CBCL Attention Problems | -4.05 | 1.24 | -3.26 | **0.009** |
| CBCL Internalizing Problems | -0.42 | 0.75 | -0.55 | 0.822 |
| CBCL Externalizing Problems | -1.07 | 0.79 | -1.36 | 0.411 |
| **State 8** |  |  |  |  |
| Intercept | 18.32 | 11.38 | 1.61 | 0.320 |
| Age | -0.07 | 0.09 | -0.87 | 0.667 |
| Sex | -0.68 | 1.31 | -0.52 | 0.822 |
| Cognition | 0.03 | 0.04 | 0.87 | 0.667 |
| Motion | -10.76 | 2.07 | -5.20 | 3.97E-06 |
| CBCL Attention Problems | 0.14 | 0.26 | 0.53 | 0.822 |
| CBCL Internalizing Problems | -0.22 | 0.16 | -1.42 | 0.402 |
| CBCL Externalizing Problems | -0.19 | 0.16 | -1.12 | 0.549 |
| **State 9** |  |  |  |  |
| Intercept | 0.63 | 0.24 | 2.60 | 0.051 |
| Age | 0.00 | 0.00 | 0.93 | 0.647 |
| Sex | 0.01 | 0.03 | 0.46 | 0.835 |
| Cognition | 0.00 | 0.00 | -0.11 | 0.991 |
| Motion | -0.20 | 0.04 | -4.48 | 1.03E-04 |
| CBCL Attention Problems | 0.00 | 0.01 | -0.66 | 0.788 |
| CBCL Internalizing Problems | 0.01 | 0.00 | 2.19 | 0.126 |
| CBCL Externalizing Problems | 0.00 | 0.00 | -0.84 | 0.691 |
| **State 10** |  |  |  |  |
| Intercept | -9.89 | 9.75 | -1.01 | 0.609 |
| Age | 0.07 | 0.07 | 0.92 | 0.653 |
| Sex | 1.31 | 1.12 | 1.17 | 0.544 |
| Cognition | 0.09 | 0.03 | 2.75 | 0.039 |
| Motion | -8.16 | 1.77 | -4.60 | 6.72E-05 |
| CBCL Attention Problems | 0.05 | 0.22 | 0.24 | 0.934 |
| CBCL Internalizing Problems | -0.22 | 0.13 | -1.62 | 0.320 |
| CBCL Externalizing Problems | -0.06 | 0.14 | -0.45 | 0.835 |
| **State 11** |  |  |  |  |
| Intercept | 21.23 | 18.94 | 1.12 | 0.549 |
| Age | 0.04 | 0.14 | 0.31 | 0.894 |
| Sex | -1.42 | 2.18 | -0.65 | 0.788 |
| Cognition | -0.01 | 0.06 | -0.08 | 0.991 |
| Motion | -21.79 | 3.44 | -6.33 | 7.78E-09 |
| CBCL Attention Problems | -0.08 | 0.43 | -0.18 | 0.959 |
| CBCL Internalizing Problems | 0.13 | 0.26 | 0.51 | 0.822 |
| CBCL Externalizing Problems | -0.39 | 0.27 | -1.43 | 0.402 |
| **State 12** |  |  |  |  |
| Intercept | 103.72 | 25.41 | 4.08 | 0.001 |
| Age | -0.31 | 0.19 | -1.60 | 0.320 |
| Sex | -2.26 | 2.93 | -0.77 | 0.719 |
| Cognition | -0.05 | 0.09 | -0.61 | 0.810 |
| Motion | -18.18 | 4.62 | -3.93 | 0.001 |
| CBCL Attention Problems | -1.20 | 0.58 | -2.07 | 0.149 |
| CBCL Internalizing Problems | -0.19 | 0.35 | -0.56 | 0.822 |
| CBCL Externalizing Problems | -0.64 | 0.37 | -1.75 | 0.265 |

Note: CBCL, Child Behavior Checklist. General Cognition is measured by the NIH Toolbox age corrected scores (28).

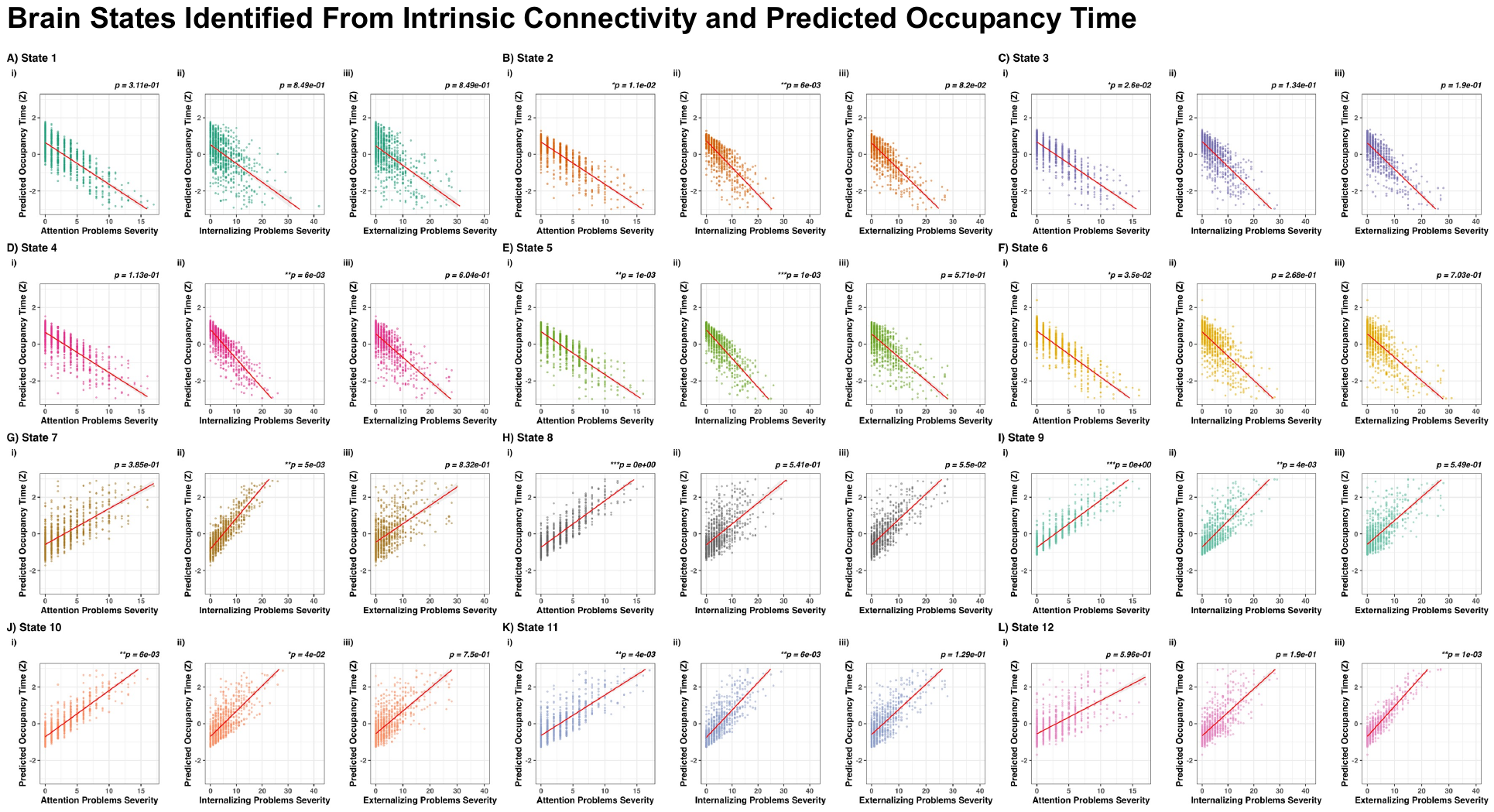

**Figure S1.** Brain states predicted occupancy time and associations with transdiagnostic symptom domains. Regression plots depict associations between predicted occupancy time (z-score) for each of the 12 brain states and severity of transdiagnostic symptom domains. P-values are FDR-corrected across all tests.

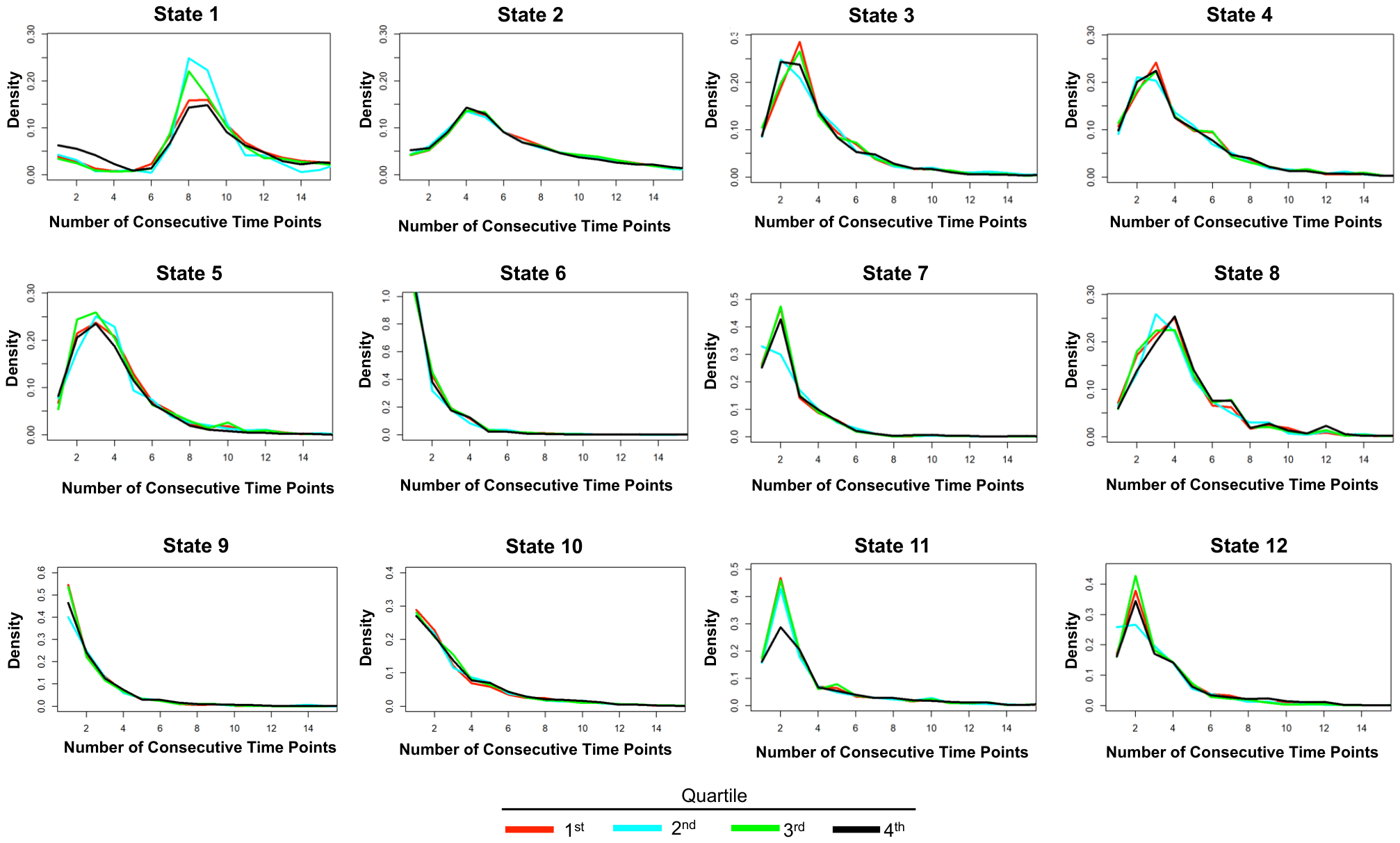

**Figure S2.** Estimated empirical sojourn distributions for each of the 12 states. Lines denote the quartile of the sojourn distribution (red=1^st^ quartile; blue=2^nd^ quartile; green=3^rd^ quartile; black=4^th^ quartile). The sojourn time represents how long a participant spends in a particular state prior to transitioning to another state.

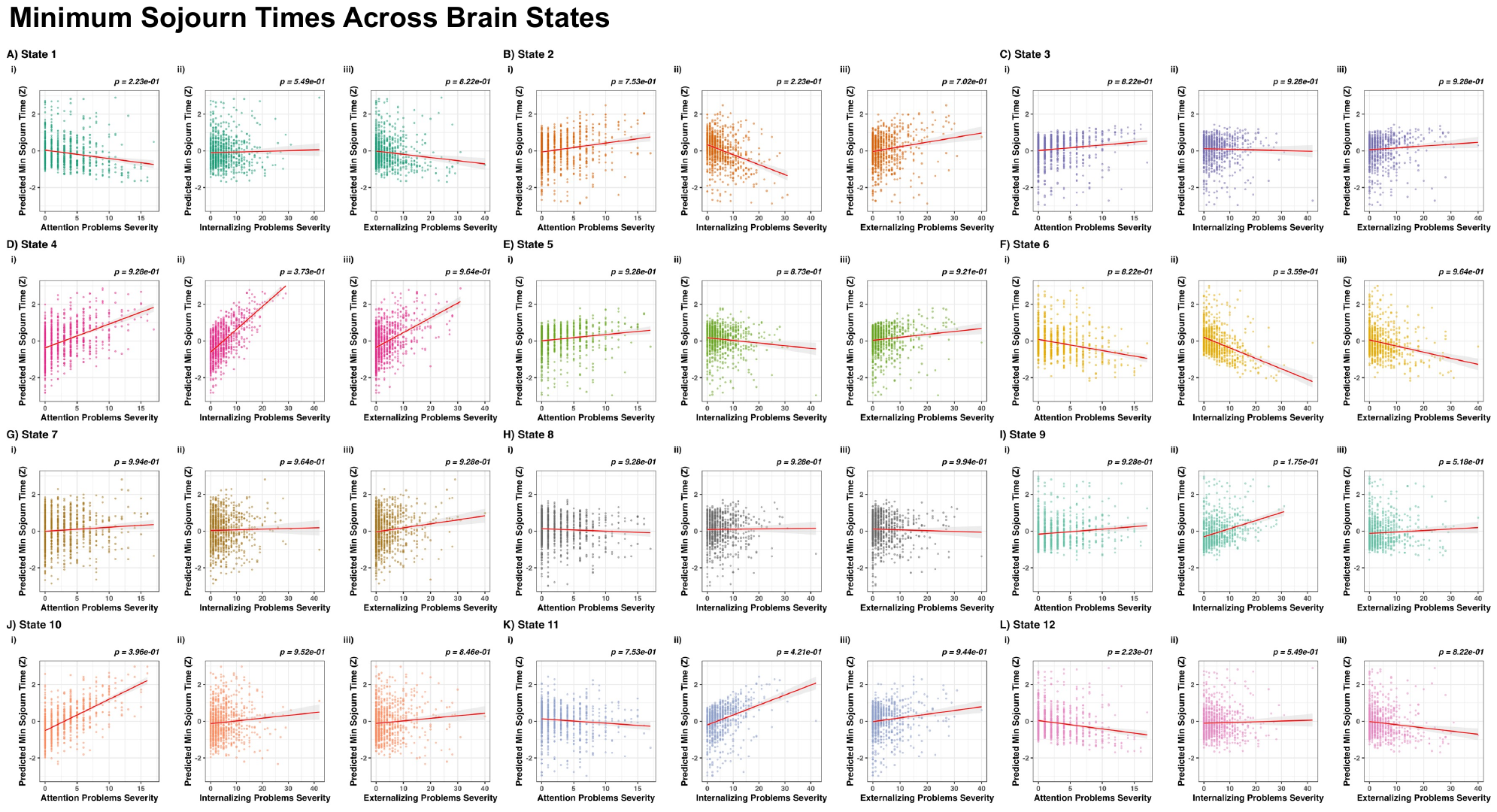
**Figure S3.** Associations between minimum sojourn time and transdiagnostic symptom domains. Regression plots depict associations between predicted minimum sojourn time (z-score) for each of the 12 brain states and severity of transdiagnostic symptom domains. P-values are FDR-corrected across all tests.

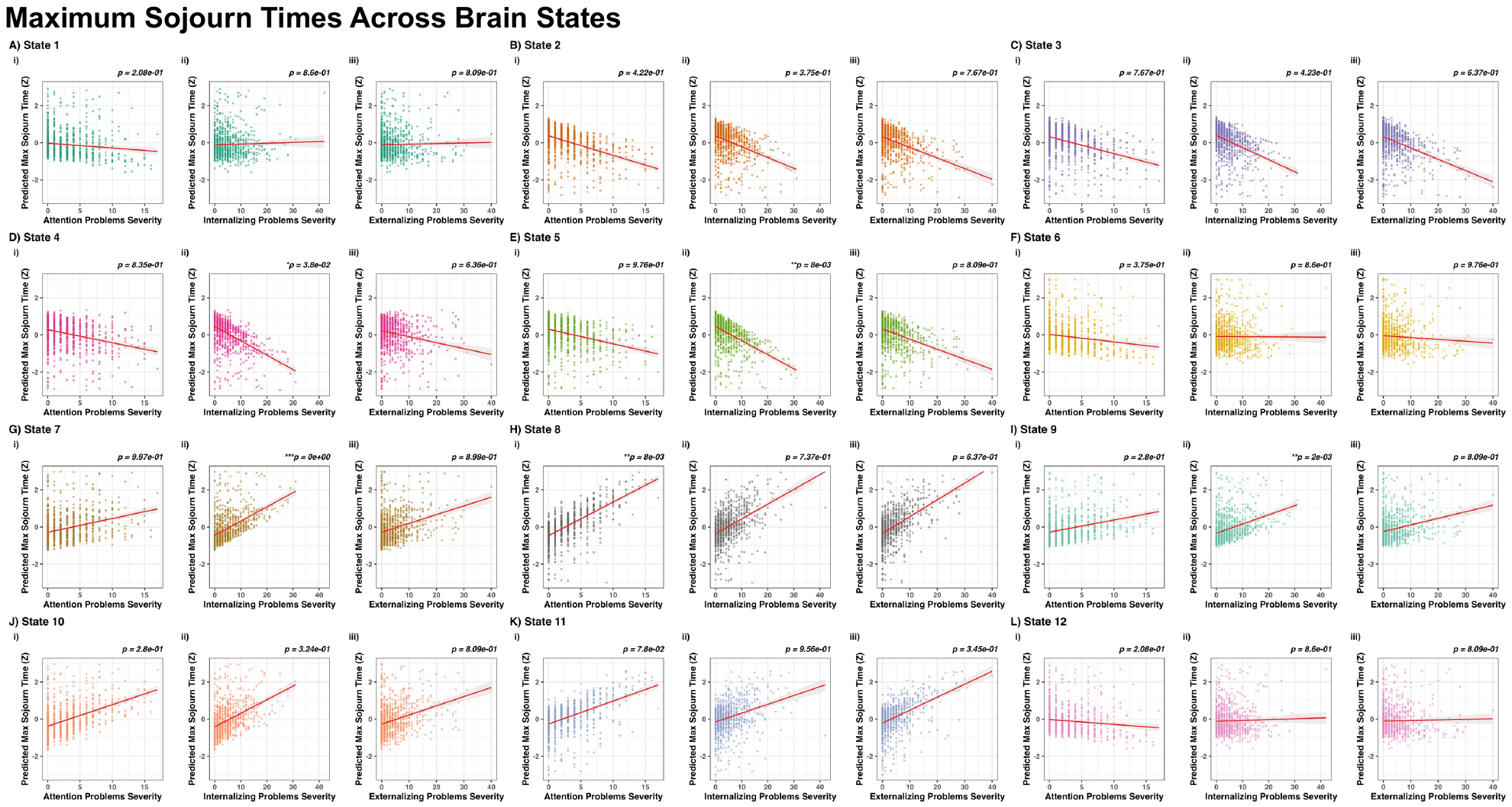
**Figure S4.** Associations between maximum sojourn time and transdiagnostic symptom domains. Regression plots depict associations between predicted maximum sojourn time (z-score) for each of the 12 brain states and severity of transdiagnostic symptom domains. P-values are FDR-corrected across all tests.

**Figure S5.** Maps of independent components and grouping into intrinsic connectivity networks for two held-out runs of resting-state. Components were derived using Independent Component Analysis (ICA) implemented using FSL MELODIC. Thirty-three independent components were mapped onto 10 large-scale networks based on the Yeo parcellation.

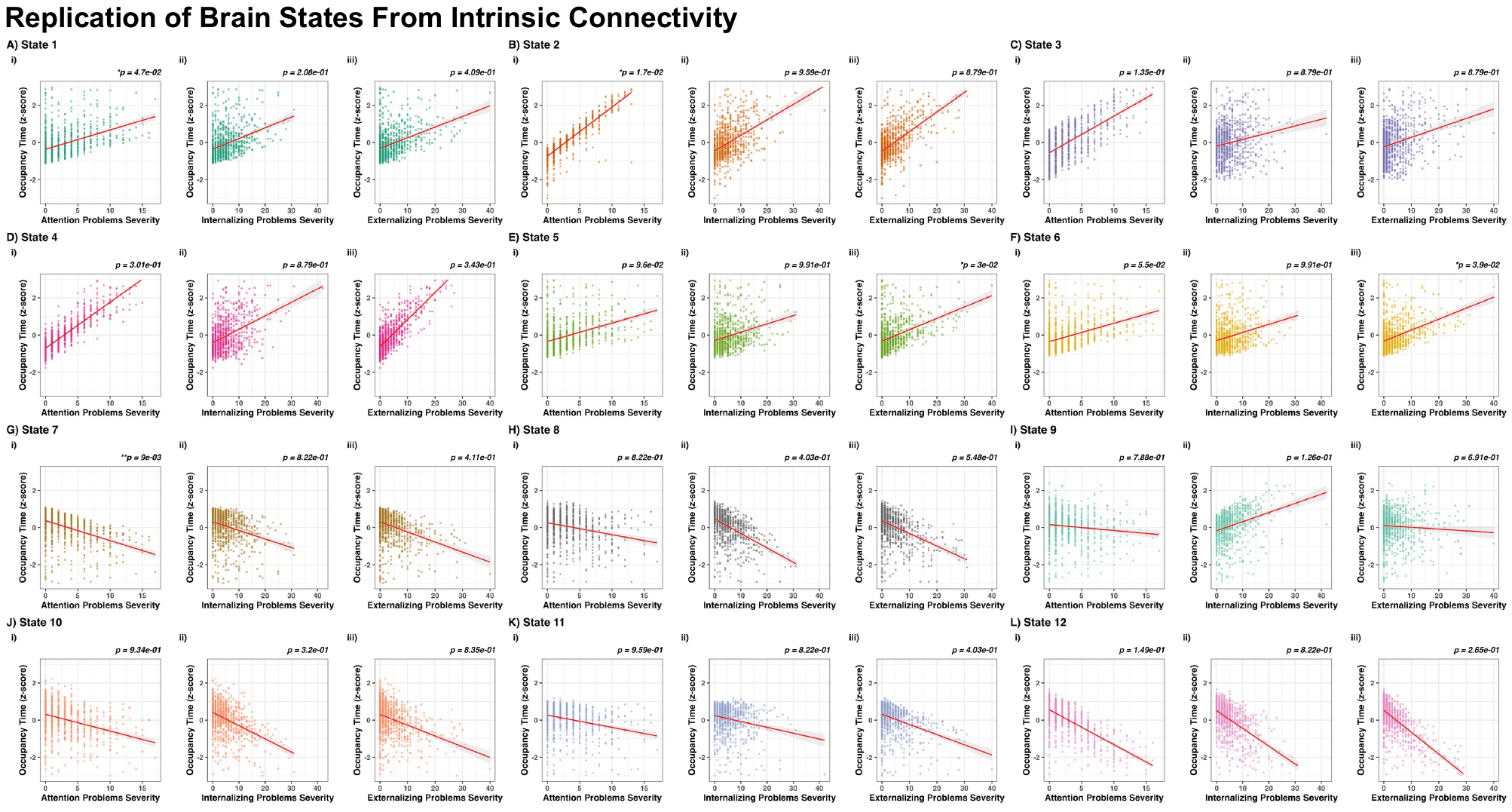
**Figure S6.** Replication of brain states predicted occupancy time and associations with transdiagnostic symptom domains. Analyses were repeated using the held-out two runs of resting-state for each participant (see main text for details). Regression plots depict associations between predicted occupancy time (z-score) for each of the 12 brain states and severity of transdiagnostic symptom domains. P-values are FDR-corrected across all tests.

**Figure S7.** Difference matrices of latent brain states compared to states 5 and 6 for the replication analysis. Given that primary analyses showed a link between occupancy time in state 12 and disruptive behavior severity, replication analyses were conducted in two, held-out resting-state runs. Heat maps present the difference in correlation values for each region pair between state 5 or 6 vs all other states. Panels Ai and Bi show correlation differences for the subset of ROI pairs with a positive correlation with state 5 and state 6, respectively, as well within the state being compared to states 5 and 6. Red cells indicate a stronger positive correlation compared to state 5 and state 6. Blue cells indicate a weaker positive correlation as compared to state 5 and state 6. White cells indicate ROI pairs with no difference between the two states. Panels Aii and Bii show correlation differences for the subset of ROI pairs with a negative correlation with state 5 and state 6, respectively, as well as within the state being compared to states 5 and 6. Red cells indicate a weaker negative correlation compared to state 5 and state 6. Blue cells indicate a stronger negative correlation as compared to state 5 and state 6. White cells indicate ROI pairs with no difference between the two states. Abbreviations for intrinsic networks: Cerblm, cerebellar; Control, Frontoparietal control network; Default, default mode network; DorsAttn, Dorsal Attention Network; Limb, limbic; Visual, visual network; SomMat, somatomotor; SC, subcortical; TP, temporo parietal; VentAttn, Ventral Attention Network. Topological representations of the top quartile of edges for all replication brain states (C). For each plot, edge widths are proportional to correlation value (i.e., darker and wider edges indicate greater positive correlations). Vertex color is specific to each intrinsic network.

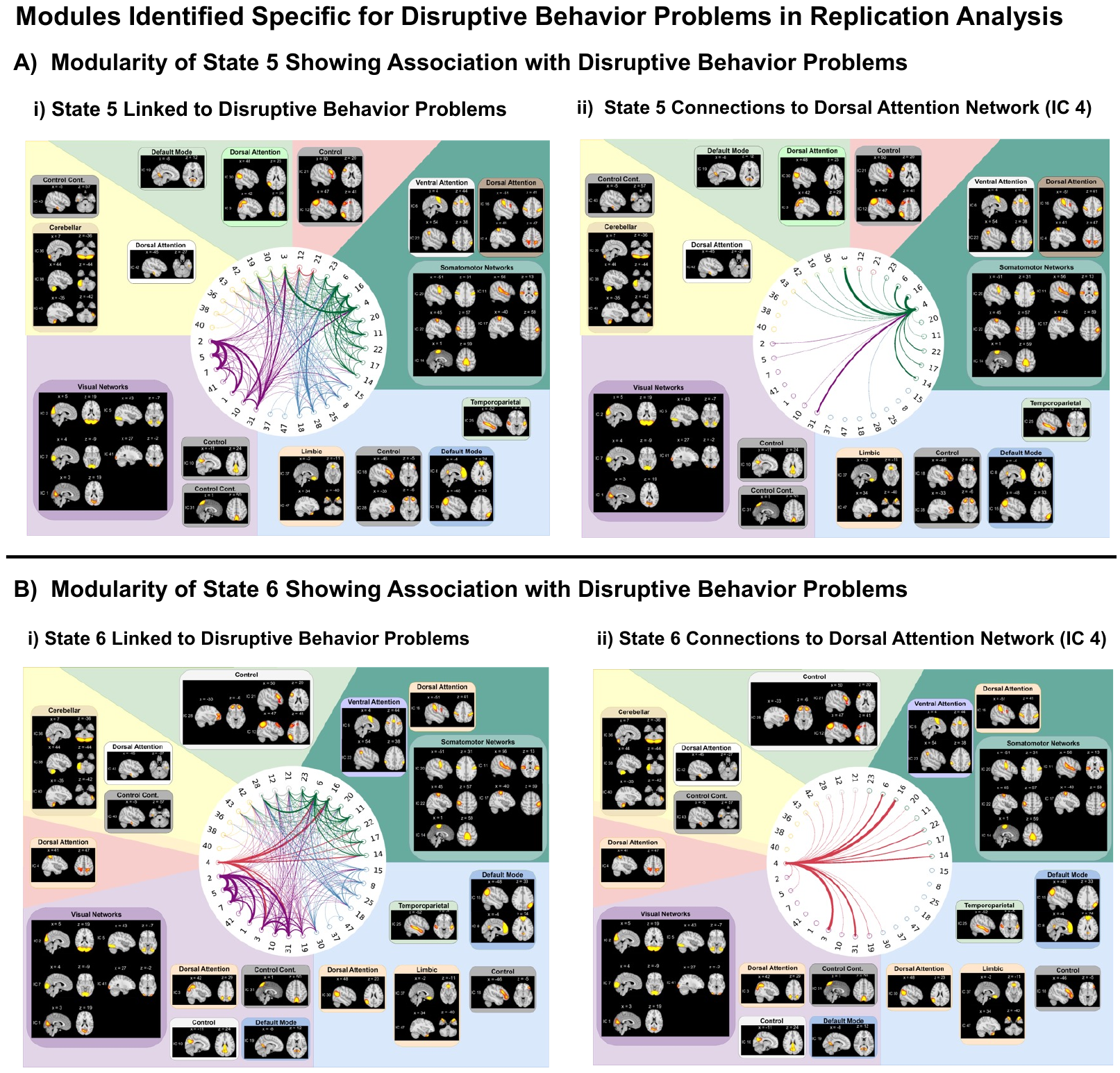
**Figure S8.** Follow-up supplemental analysis of modularity for replication states specific to disruptive behavior problems. In replication analyses in resting-state runs 3-4, for any states emerging that showed significant and distinct or specific associations with disruptive behavior problems, we examined modularity to further understand state characteristics. Thus, we used a modularity analysis to identify community structures within the state. State 5 (Ai) and state 6 (Bi) were significantly and distinctly associated with disruptive behavior problems in replication analyses, and we conducted modularity analyses. IC 4 formed a single, distinct module in state 6, indicating that its connection patterns in state 6 were markedly different from those in state 5. Although IC 4 does not include the identical regions as IC 24 in our primary analysis, it is also part of the dorsal attention network, further emphasizing changes within this network associated with states linked to disruptive behavior.

**Supplemental Discussion:**

Follow-up modularity analyses indicated potential perturbations within the dorsal attention network in brain states associated with disruptive behavior problems. Specifically, state 12 showed weaker connections between the dorsal attention and somatomotor networks. Additionally, state 12 showed weaker connections within somatomotor circuitry and a less distributed pattern of connections between the dorsal attention network and other networks (i.e., in state 12, there were fewer connections between the dorsal attention network IC 24 to all other modules vs state 2). Prior work has shown that externalizing-related behaviors and symptom domains is associated with aberrant patterns of dynamic functional connectivity within cognitive control networks, including the dorsal attention network (32, 33). The dorsal attention network is a sub-circuitry of cognitive control networks and is implicated in attentional processes including working memory (34-36). The dorsal attention network also has distributed functional connections with related cognitive control sub-networks including the frontoparietal, ventral attention, and default mode networks (37). Altered activity and connectivity within the dorsal attention network has also been shown to be related to disruptive behavior problems (38-40). Additionally, cognitive control sub-networks, including dorsal/ventral attention, frontoparietal, and default mode networks, are among the most globally connected in the brain and are characterized as core hubs of integration for cognitive control and social perception processes (41, 42). Results of follow-up analyses for modularity indicated a less distributed pattern of connections among the dorsal attention network in state 12 and could suggest reduced segregation of cognitive control sub-networks, which aligns with prior findings reporting reduced modularity and segregation of networks in ADHD (10), schizophrenia (43), and ASD (44).
